# MetaCurator: A hidden Markov model-based toolkit for extracting and curating sequences from taxonomically-informative genetic markers

**DOI:** 10.1101/672782

**Authors:** Rodney T. Richardson, Douglas B. Sponsler, Harper McMinn-Sauder, Reed M. Johnson

## Abstract

The community-level analysis of samples containing diverse genetic material, via metabarcoding and metagenomic approaches, is increasingly popular. While the production of sequence data for such studies has become straightforward, questions remain about how best to analyze and taxonomically characterize sequence data. For many sequence classification approaches, an important component of the workflow involves the curation of reference sequences. Ideally, this involves trimming away extraneous sequence at the 3 prime and 5 prime ends of the target marker of interest, as well as the removal of reference sequence duplicates. Here, we present MetaCurator, a software package written in Python, designed for automated reference sequence curation and highly generalizable across markers and study systems. MetaCurator is organized in a modular fashion, so users can implement tools individually in addition to utilizing the automated and flexible MetaCurator parental code. Aside from modules used to organize and format taxonomic lineage data, MetaCurator contains two signature tools. IterRazor utilizes profile hidden Markov models and an iterative search framework to exhaustively identify and extract the precise amplicon marker of interest from available reference sequence data. DerepByTaxonomy then facilitates sequence dereplication using a taxonomically aware approach, removing duplicates only when they belong to the same taxon. This is important for cases of incomplete lineage sorting between species and for highly conserved markers, such as plant *rbcL* and *trnL*, which often display no sequence divergence across taxa, even at the genus level.

**Availability and implementation:** MetaCurator is supported on OSX and Linux (RedHat/CentOS) and is freely available under a GPL v3.0 license at https://github.com/RTRichar/MetaCurator.

**Supplemental information:** Code associated with this work is available at https://github.com/RTRichar/MetabarcodeDBsV2 and additional analysis is presented in supplementary files.

## 1 Introduction

During sequence-based characterization of biological communities, the supervised taxonomic classification of sequences is an important goal. Numerous sequence classification software programs accomplish this by measuring sequence similarity and modelling relationships between sequence similarity and taxonomic affiliation. Such classifiers often rely upon strong, and frequently violated, assumptions concerning database curation.

Approaches to identifying and extracting marker sequence data from disparate sources exist but are typically designed for specific markers or taxa (Bengtsson-Palme et al. 2013; Ankenbrand et al. 2015), with the exception of ANACAPA and the Metaxa2 Database Builder (Bengtsson-Palme et al. 2018; Curd et al. 2019). Additionally, *in silico* PCR techniques exist for analyzing marker sequences (Ficetola et al. 2010; Elbrecht and Leese, 2017), though these pipelines are intended to improve metabarcoding primer universality as opposed to aiding database curation.

The use of poorly curated reference databases and taxonomically naïve sequence dereplication methods are important methodological shortcomings for metabarcoding. During composition-based classification, extraneous sequence regions adjacent to the marker of interest bias the *k*-mer distributions used by many tools, such as SINTAX, the RDP Naïve Bayesian Classifier and VSEARCH (Wang et al. 2007; Edgar 2016; Rognes et al. 2017). For classifiers which estimate taxonomic boundaries by analyzing the distance distributions of the top *N* aligned sequences (Huson et al. 2011; Bengtsson-Palme et al. 2015; Gao et al. 2017), retention of sequence duplicates can bias characterization of the distance distribution between query sequences and top hit reference taxon matches, especially as the number of top hit taxon replicates approaches *N*. However, taxonomically naïve removal of sequence duplicates without ensuring that the two sequences are from the same taxon represents the removal of important information from the database regarding the lack of genetic variation between two taxa.

For these issues, we developed the MetaCurator toolkit. The main tool, IterRazor, identifies and extracts target marker reference sequences from any available source, including whole genome assemblies for example. After extracting the references for a given marker, DerepByTaxonomy is used to dereplicate reference sequences using a taxonomy-aware approach. Additional Python tools and a Unix shell script facilitate the formatting of taxonomic lineages for hierarchical curation and the removal of taxonomic lineage artifacts commonly found within the NCBI Nucleotide Database (NCBI Resource Coordinators, 2018).

## 2 Features

### 2.1 IterRazor target amplicon extraction algorithm

Using MAFFT (Katoh and Standley, 2013), the IterRazor algorithm produces a seed multiple sequence alignment (MSA) of approximately 5 to 10 reference sequences trimmed to the exact marker of interest and provided by the user. This MSA is used as input for the hmmbuild algorithm of HMMER3 (Eddy, 2011), producing an initial HMM profile of the marker. After preparing the HMM profile for searching using hmmpress, the algorithm proceeds into a multi-round, multi-iteration search. Heterogeneous input query sequences (e.g. whole chloroplast genomes or sanger sequences) from diverse taxa, are searched against this profile using nhmmscan. Tabular HMM search output is parsed for profile matches which meet the user-specified quality standards. Sequences exhibiting a sufficient match to the HMM profile are trimmed to the matching region and aligned individually to the existing MSA of marker sequences using hmmalign. An updated HMM profile is then calculated with the new alignment using hmmbuild. Following each iteration, sequences from which the marker was extracted are removed from the query sequences to decrease extraction time. With this approach, the corpus of references can increase with each iteration and the HMM profile becomes more representative of the marker and taxonomic group of interest as sequences from novel taxa are incorporated. To avoid the addition of false positive HMM matches, users can adjust the HMM profile match E-value threshold using the ‘--HmmEvalue’ argument, set to a default of 0.005.

The user can define the number of extraction rounds, iterations per round and the percent profile match coverage of the HMM profile required for each round. By default, IterRazor conducts four rounds of extraction, requiring profile matches to cover 100, 95, 90 and 65 percent of the HMM profile per round, with 20, 10, 5 and 5 search iterations per round. If a search iteration fails to yield new extracted sequences, IterRazor will break out of the iteration loop and proceed to the subsequent search round. After the final iteration of the final round, an output file containing all extracted sequences is written and the number of sequences extracted in each iteration is printed to standard output. Additionally, a temporary directory containing the intermediary files produced during extraction is deleted unless ‘--SaveTemp True’ is declared. Default IterRazor settings work well for high-conservation markers. Low-conservation, length-variable markers sometimes require more stringent HMM search E-value and percent profile match coverage thresholds.

### 2.2 Sequence dereplication by taxonomy

Provided with a fasta file of reference sequences and a text file containing associated taxonomic lineages, DerepByTaxonomy utilizes VSEARCH-based alignment (Rognes et al. 2017) to find exact match semi-global alignments, test if the associated lineages belong to the same taxon and, if so, remove the shorter of the two duplicates. To accomplish this, DerepByTaxonomy iteratively searches the fasta sequence corpus against itself requiring 100 percent identity matches and 100 percent query coverage while calculating alignments using an infinite internal gap penalty and a null external gap penalty. By default, DerepByTaxonomy performs 10 search rounds during dereplication. Since VSEARCH uses *k*-mer composition to prioritize a limited number of alignment tests (set by ‘--maxaccepts’ and ‘--maxrejects’), multiple dereplication rounds ensure sequences will be exhaustively searched for duplicates. If no duplicates are found for a search round, the tool breaks from the loop and subsequent rounds are not performed.

### 2.3 Automated database curation with MetaCurator parental module

In addition to implementing the MetaCurator tools separately, users can execute the entire pipeline using the MetaCurator command. This parental code runs taxonomy reformatting, cleaning and mid-point correction; IterRazor marker sequence extraction; and DerepByTaxonomy dereplication while allowing the same options flexibility as implementing each sub-component individually.

## 3 Results

Using MetaCurator, we constructed databases for five metabarcoding markers (Table 1), including the low-conservation plant *trnH* and ITS2 markers. The primer sets used to designate these barcode regions are detailed in Supplemental Table S1. Relative to previous studies (Richardson et al. 2018 and 2019), between 14.3 to 50.7 percent more genera were retained during data curation. Further, while 1.8 to 34.1 percent of genera were lost during database dereplication in these previous studies, no taxa were lost during MetaCurator dereplication.

**Table 1:**
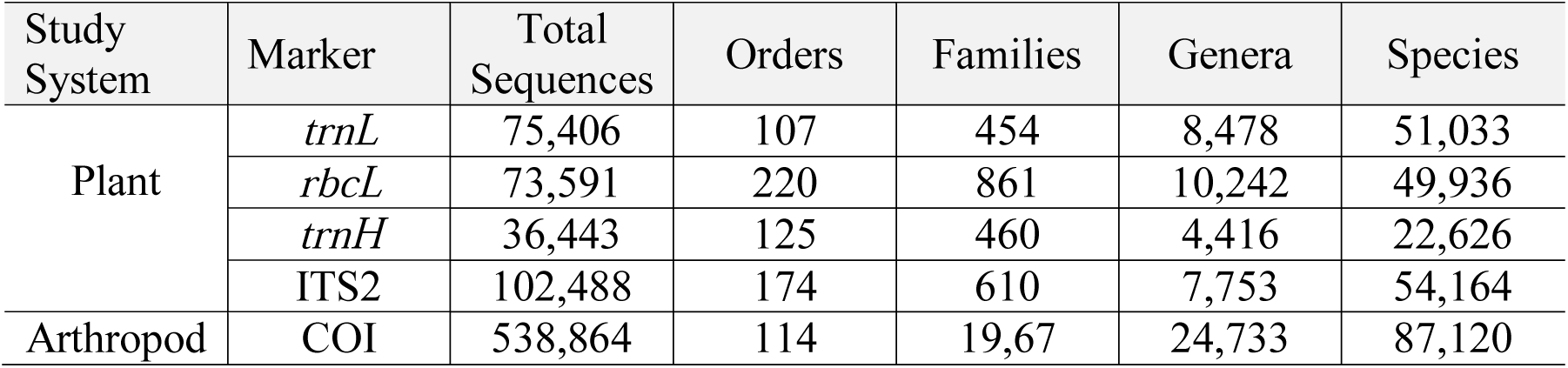
Summary of the taxonomic richness of each database produced in this study.

| Study System | Marker | Total Sequences | Orders | Families | Genera | Species |
| --- | --- | --- | --- | --- | --- | --- |
| Plant | *trnL* | 75,406 | 107 | 454 | 8,478 | 51,033 |
|  | *rbcL* | 73,591 | 220 | 861 | 10,242 | 49,936 |
|  | *trnH* | 36,443 | 125 | 460 | 4,416 | 22,626 |
|  | ITS2 | 102,488 | 174 | 610 | 7,753 | 54,164 |
| Arthropod | COI | 538,864 | 114 | 19,67 | 24,733 | 87,120 |

Visualizing the number of sequences added per iteration and per round revealed that large fractions of sequences were extracted after the initial iteration, particularly for low-conservation markers (Supplemental Figure S1). In testing MetaCurator on *rbcL* and ITS2 data using a fixed profile HMM length coverage threshold of 85 percent, the resulting sequence length distributions were shorter relative to the recommended stepwise length coverage thresholds (two-tailed Wilcoxon rank sum test, *P* < 0.0001 for both markers). In analyzing multiple sequence alignments of 50 random marker sequences from ITS2 and *rbcL*, both alignments produced regions of high consensus across all sequences (Supplemental Figures S2 and S3), suggesting a low rate of false discoveries during extraction. Further characterization is provided in the Supplemental Material.

## Supporting information

MetaCurator Manual

## Funding

This work was supported by a Project Apis m. – Costco Honey Bee Biology Fellowship to RTR and support provided by state and federal funds appropriated to The Ohio State University, Ohio Agricultural Research and Development Center.

## Notes

https://github.com/RTRichar/MetaCurator/tree/master/TestMetaCurator

https://github.com/RTRichar/MetabarcodeDBsV2

## References

Ankenbrand MJ, Keller A, Wolf M, Schultz J, Förster F (2015) ITS2 database V: Twice as much. Molecular Biology and Evolution, 32, 3030–3032.

Bengtsson-Palme, J, M Ryberg, M Hartmann, et al. (2013) Improved software detection and extraction of ITS1 andITS2 from ribosomal ITS sequences of fungi and other eukaryotes for analysis of environmental sequencing data. Methods in Ecology and Evolution, 4, 914–919.

Bengtsson-Palme, J, M Hartmann, KM Eriksson, C Pal, K Thorell, DGJ Larsson, RH Nilsson (2015) Metaxa2: improved identification and taxonomic classification of small and large subunit rRNA in metagenomic data. Molecular Ecology Resources, 15, 1403–1414.

Bengtsson-Palme, J, RT Richardson, M Meola, et al. (2018) Metaxa2 Database Builder: enabling taxonomic identification from metagenomic or metabarcoding data using any genetic marker. Bioinformatics, 34, 4027–4033.

Curd, EE, Z Gold, G Kandlikar, et al. (2019) Anacapa Toolkit: An environmental DNA toolkit for processing multilocus metabarcode datasets. Methods in Ecology and Evolution, https://doi.org/10.1111/2041-210X.13214.

Eddy, SR (2011) Accelerated Profile HMM Searches. PLoS Computational Biology, 7, e1002195.

Edgar RC (2016) SINTAX: A simple non-Bayesian taxonomy classifier for 16S and ITS sequences. Biorxiv, 074161.

Elbrecht, V, F Leese (2017) PrimerMiner: an r package for development and in silico validation of DNA metabarcoding primers. Methods in Ecology and Evolution, 8, 622–626.

Ficetola, GF, E Coissac, S Zundel, T Riaz, W Shehzad, J Bessiere, P Taberlet, F Pompanon (2010) An *in silico* approach for the evaluation of DNA barcodes. BMC Genomics, 11, 434.

Gao, X, H Lin, K Revanna, Q Dong (2017) A Bayesian taxonomic classification method for 16S rRNA gene sequences with improved species-level accuracy. BMC Bioinformatics, 18, 247.

Huson, DH, S Mitra, H-J Ruscheweyh, N Weber, SC Schuster (2011) Integrative analysis of environmental sequences using MEGAN 4. Genome Research, 21, 1552–1560.

Katoh, K, Standley DM (2013) MAFFT multiple sequence alignment software version 7: improvements in performance and usability. Molecular Biology and Evolution, 30, 772–780.

NCBI Resource Coordinators (2018) Database resources of the national center for biotechnology information. Nucleic Acids Research, 46, D8–D13.

Ohio Supercomputer Center. 1987. Ohio Supercomputer Center. Columbus OH: Ohio Supercomputer Center. http://osc.edu/ark:/19495/f5s1ph73.

Richardson, RT, J Bengtsson-Palme, MM Gardiner & RM Johnson. (2018) A reference cytochrome c oxidase subunit I database curated for hierarchical classification of arthropod metabarcoding data. PeerJ, 6, e5126.

Richardson, RT, HR Curtis, C-H Lin, DB Sponsler & RM Johnson. (2019). Honey bee spring foraging patterns in Midwestern corn and soy agroecosystems revealed with quantitative metabarcoding and waggle dance inference. Molecular Ecology, 28, 686–697.

Rognes, T, T Flouri, B Nichols, C Quince, F Mahé (2017) VSEARCH: a versatile open source tool for metagenomics. PeerJ, 4, e2584.

Wang, Q, GM Garrity, JM Tiedje, JR Cole (2007) Naïve Bayesian classifier for rapid assignment of rRNA sequences into the new bacterial taxonomy. Applied and Environmental Microbiology, 73, 5261–5267.

