## Supplementary material for "MetaCurator: A hidden Markov model-based toolkit for extracting and curating sequences from taxonomically-informative genetic markers": MetaCurator Manual

**Users Guide**

MetaCurator: Curation Toolkit for Barcoding, Metabarcoding and Metagenetic Reference Sequence Databases

### 1. Conceptual visualization

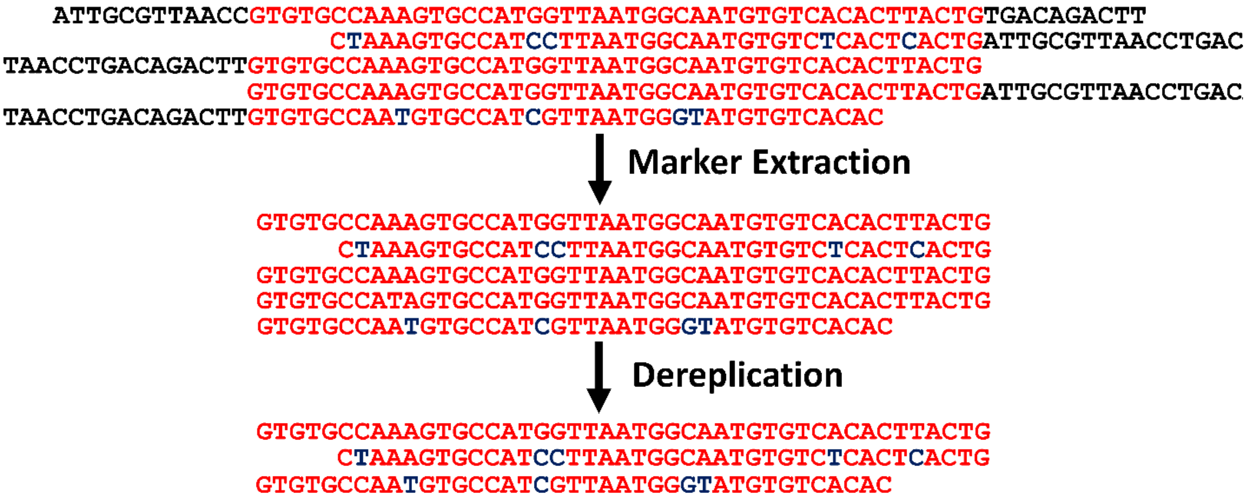


### 1. Introduction

This documentation guides users through the acquisition, installation and use of the MetaCurator curation toolkit, which can be found at https://github.com/RTRichar/MetaCurator. The major functionality of this software is encoded in an algorithm, IterRazor, designed to identify and extract the barcode region of interest from diverse sources of reference sequence data, including whole genomes or whole mitochondrion genomes. Having a reference database representative of a particular marker of interest is important for the taxonomic model selection procedures of various DNA sequence classifiers. Further, it improves the researcher's ability to accurately inventory the taxa represented in the database with respect to specific markers used in barcoding, metabarcoding and metagenomics.

This package also includes software for dereplicating extracted sequences using a taxonomically-aware method, DerepByTaxonomy – dereplication by taxonomy – which only removes sequence duplicates if the two duplicate sequences belong to the same taxon. For many barcode markers, such as plant *rbcL*, taxonomically-unsupervised dereplication is a common oversight as many identical sequences belong to different species, and even different genera in many cases. In the context of DNA barcoding and metagenetic analysis, a legitimate lack of genetic divergence between taxa at a particular locus is important to retain in the reference database in order to decrease the probability of sequence overclassification to high-resolution ranks, such as species and genus.

While the IterRazor extraction software and DerepByTaxonomy dereplication software are available as separate tools along with a number of helper formatting scripts, 'MetaCurator.py' is provided as a parental module which streamlines the recommended curation process into a single command (skip to section 3.4 for details). All python tools provided are encoded with a help screen, which can be accessed by declaring '-h' or '--help.'

### 2. Installation

#### *2.1 Dependencies*

- Python 2 or Python 3
- Perl 5.16.0 or compatible
- VSEARCH 2.8.1 or compatible
- HMMER 3.1b2 or compatible
- MAFFT 7.270 or compatible

#### *2.2 Environmental Configuration*

After unpacking MetaCurator_v1.0.0.tar.gz, export the path of the directory containing the executable python files to $PATH and ensure that all files with '.py' and '.sh' have executable privileges (run 'chmod 755 file.py' if not). If using an ssh client on a cluster environment, add an export command to the appropriate bash profile file. Further, ensure that VSEARCH, MAFFT and HMMER dependencies are added to $PATH.

### 3. Algorithms

#### *3.1 IterRazor.py*

```
IterRazor.py -r {InputReferences.fasta} -i {Input.fasta} -it {Input.tax} -o {Output.fasta}
```

This program identifies and excises the precise barcode marker of interest from the provided input reference sequences.

Input formatting:

For '-r', IterRazor requires an input fasta file containing 5 to 10 manually trimmed sequences, formatted as below, representative of a broad diversity of the taxa of interest (e.g. for seed plants, one might include representatives from Ginkgoales, Pinales, Magnoliales, Poales, Apiales, Vitales, Rosales and Brassicales). These sequences must be trimmed to the exact barcode marker region of interest as they will be used to construct the seed profile HMM for homology searching.

```
>MG222568.1
TGTGTGGACCGATGGACTTACTAGCCTTGATCGTTACAAAGGGCGATGCTACCACATCGAGCCCGTTGCTGGAGAAGAAAATCAATTTATT
GCTTATGTAGCTTATCCTTTAGACCTTTTTGAAGAAGGGTCTGTTACTAATATGTTTACTTCCATTGTGGGTAATGTATTTGGGTTCAAAGCG
```

For ‘-i’, IterRazor requires an input fasta file of reference sequences to be trimmed, typically with NCBI accessions as headers (fasta headers have to be compatible with taxonomy headers as discussed below).

```
>MG222568.1
CAGCTTTCCGAGTAACTCCTCAACCTGGGGTTCCACCTGAAGAAGCAGGGGCCGCGGTAGCTGCCGAATCTTCTACTGGTACATGGACAAC
TGTGTGGACCGATGGACTTACTAGCCTTGATCGTTACAAAGGGCGATGCTACCACATCGAGCCCGTTGCTGGAGAAGAAAATCAATTTATT
GCTTATGTAGCTTATCCTTTAGACCTTTTTGAAGAAGGGTCTGTTACTAATATGTTTACTTCCATTGTGGGTAATGTATTTGGGTTCAAAGCG
CTGCGCGCTCTACGTCTGGAAGATCTGCGAATCCCTACTGCGTATGTTAAAACTTTCCAAGGACCGCCTCATGGTATCCAAGTTGAAAGAGA
TAAATTGAACAAGTATGGTCGTCCCCTGTTGGGATGTACTATTAAACCTAAATTGGGGTTATCTGCTAAAAACTACGGTAGAGCAGTTTATG
AATG
```

For ‘-it’, IterRazor requires an input taxonomy file containing lineages associated with sequences, typically in Metaxa2-compatible format (as long as fasta and taxonomy file headers are compatible and taxonomy headers are tab-separated from lineages, lineages can be custom formatted). For certain classifiers, such as Metaxa2, higher resolution ranks can be omitted for cases of ambiguous open nomenclature (e.g. accession KF284759.1 below is not identified to species). The curation toolkit is made to accommodate this but such entries may need to be removed for alternative classifiers such as SINTAX or RDP.

```
NC_035751.1	k__Eukaryota;p__Streptophyta;c__Liliopsida;o__Poales;f__Poaceae;g__Miscanthus;s__Miscanthus junceus;
EU697437.1	k__Eukaryota;p__Streptophyta;c__Jungermanniopsida;o__Metzgeriales;f__Aneuraceae;g__Aneura;
KF284759.1	k__Eukaryota;p__Streptophyta;c__Liliopsida;o__Alismatales;f__Araceae;g__Colocasia;
```

#### *3.2 DerepByTaxonomy.py*

*DerepByTaxonomy.py -i {Input.fasta} -t {Input.tax} -o {Output.fasta}*

This program uses a VSEARCH-based approach to remove sequence duplicates, sequences which are either globally identical or semi-globally identical to a subsection of another sequence. A duplicate sequence is only removed if both sequences have the same taxonomic annotations.

Input formatting:

For ‘-i’, DerepByTaxonomy requires an input fasta file of reference sequences, typically with NCBI accessions as headers.

```
>MG222568.1
CAGCTTTCCGAGTAACTCCTCAACCTGGGGTTCCACCTGAAGAAGCAGGGGCCGCGGTAGCTGCCGAATCTTCTACTGGTACATGGACAAC
TGTGTGGACCGATGGACTTACTAGCCTTGATCGTTACAAAGGGCGATGCTACCACATCGAGCCCGTTGCTGGAGAAGAAAATCAATTTATT
GCTTATGTAGCTTATCCTTTAGACCTTTTTGAAGAAGGGTCTGTTACTAATATGTTTACTTCCATTGTGGGTAATGTATTTGGGTTCAAAGCG
CTGCGCGCTCTACGTCTGGAAGATCTGCGAATCCCTACTGCGTATGTTAAAACTTTCCAAGGACCGCCTCATGGTATCCAAGTTGAAAGAGA
TAAATTGAACAAGTATGGTCGTCCCCTGTTGGGATGTACTATTAAACCTAAATTGGGGTTATCTGCTAAAAACTACGGTAGAGCAGTTTATG
AATG
```

For ‘-t’, DerepByTaxonomy requires input taxonomy file containing lineages associated with sequences, typically in Metaxa2-compatible format. The major requirement is that the tab-separated lineage headers must match those of the input fasta.

```
NC_035751.1	k__Eukaryota;p__Streptophyta;c__Liliopsida;o__Poales;f__Poaceae;g__Miscanthus;s__Miscanthus junceus;
EU697437.1	k__Eukaryota;p__Streptophyta;c__Jungermanniopsida;o__Metzgeriales;f__Aneuraceae;g__Aneura;
KF284759.1	k__Eukaryota;p__Streptophyta;c__Liliopsida;o__Alismatales;f__Araceae;g__Colocasia;
```

*3.3 Helper tools*

*Rtaxa2Mtaxa.py*: After obtaining taxonomies using Taxonomizr R function, convert to Metaxa2-usable format with this tool. This Metaxa2 format is similar to the formats required by many other DNA sequence classifiers, such as SINTAX and RDP, so users can easily reformat the lineages using Perl regular expression-based substitution for further cross-compatibility.

*Rtaxa2Mtaxa.py -i {TaxonomizrOutput.csv} -o {Output.tax}*

*CleantTax.sh*: Shell script containing Perl regular expression commands for cleaning up NCBI artifacts from lineages. The input file should be suffixed with '.tax' and the output filename will have this part of the filename replaced with '_clean.tax.'

*CleantTax.sh {Input.tax}*

*ReviseIntNAs.py*: Revise 'NA' instances, representative of unresolved taxonomy, at intermediate nodes in the taxonomic lineages.

*ReviseIntNAs.py -i {Input.tax} -o {Output.tax}*

*TaxFastaConsensus.py*: Program for removing sequences which don't have associated lineages and simultaneously removing lineages which don't have associated sequences.

*TaxFastaConsensus.py -it {Input.tax} -if {Input.fasta} -ot {Output.tax} -of {Output.fasta}*

*3.4* ***MetaCurator.py***

*MetaCurator.py -r {InputReferences.fasta} -i {Input.fasta} -it {Input.tax} -of {Output.fasta} -ot {Output.tax}*

As the parental code of the curation toolkit, MetaCurator implements the IterRazor and DerepByTaxonomy algorithms and makes use of the helper tools in a streamlined procedure. Using MetaCurator, the user provides input sequences, a taxonomy file and a set of references trimmed to the barcode region of interest. Further, the user can specify any argument from the individual extraction and dereplication algorithms. For added functionality, MetaCurator can process taxonomic lineage output directly from Taxonomizr by declaring '--TaxonomizrFormat True.'

Required arguments:

| | |
|---|---|
| -r, --ReferencesFile | Fasta file of sequences trimmed to exact region of interest (header format should conform to inputfile header format). This should be 5 to 10 sequences spanning a diversity of the phylogenetic tree of interest, so as to build an appropriately inclusive HMM from the first iteration. |
| -i, --InputFile | Input file of reference sequences to be extracted (sequence headers should include a unique sequence identifier, NCBI accessions are recommended, preceded only by '>') |

| | |
|---|---|
| -it, --InTax | Input taxonomic lineages (a tab-delimited file with the first column containing unique sequence identifiers compatible with the files provided under '-r' and '-i'). Normally, this would be in Metaxa2-compatible format, however, if lineages come directly from Taxonomizr, you can skip reformatting and declare '-tf True.' See below for more details. |
| -ot, --OutTax | Filename for output taxonomic lineages |
| -of, --OutFasta | Filename for output fasta formatted sequences |

Universal optional arguments:

| | |
|---|---|
| -t, --threads | Number of processors to use |
| -st, --SaveTemp | Option for saving alignments and HMM files produced during extraction and dereplication (True or False) |
| -tf, --TaxonomizrFormat | Specify if tax lineages are in Taxonomizr format and must first be converted to Metaxa2-compatible tab-delimited format (True or False) |
| -ct, --CleanTax | Specify if tax lineages are to be cleaned of common NCBI artifacts and revised at unresolved midpoints (True or False) |

IterRazor-specific optional arguments:

| | |
|---|---|
| -e, --HmmEvalue | Evalue threshold for HMM search |
| -is, --IterationSeries | The IterationSeries and CoverageSeries arguments dictate the number of iterative searches to run and the minimum HMM coverage required for each round of extracting. This argument consists of a comma-separated list specifying the number of searches to run during each round of extraction. The list can vary in length, changing the total number of extraction rounds conducted. However, if a search iteration fails to yield any new reference sequence amplicons, IterRazor will break out of that round of searching and move to the next. By default, the software conducts four rounds of extraction with 20, 10, 5 and 5 search iterations per round (i.e. '-is 20,10,5,5'). |
| -cs, --CoverageSeries | This argument consists of a comma-separated list specifying the minimum HMM coverage required per round for an extracted amplicon reference sequence to be added to the reference sequence fasta. The list can vary in length but must be equal in length to the IterationSeries list. By default, the software conducts four rounds of extraction with coverage limits of 1.0, |

| | |
|---|---|
| | 0.95, 0.9 and 0.85 for each respective search round (i.e. '-cs 1.0,0.95,0.9,0.85'). |

DerepByTaxonomy-specific optional arguments:

| | |
|---|---|
| -mh, --MaxHits | Max hits to find before ending search for any given query using VSEARCH (default: 1000) |
| -di, --Iterations | Number of iterative VSEARCH searches to run (default: 10). This is important in case the number of replicates for a given sequence greatly exceeds '--MaxHits' |

Input formatting:

Requires an input fasta file of reference sequences, typically with NCBI accessions as headers.

```
>MG222568.1
CAGCTTTCCGAGTAACTCCTCAACCTGGGGTTCCACCTGAAGAAGCAGGGGCCGCGGTAGCTGCCGAATCTTCTACTGGTACATGGACAAC
TGTGTGGACCGATGGACTTACTAGCCTTGATCGTTACAAAGGGCGATGCTACCACATCGAGCCCGTTGCTGGAGAAGAAAATCAATTTATT
GCTTATGTAGCTTATCCTTTAGACCTTTTTGAAGAAGGGTCTGTTACTAATATGTTTACTTCCATTGTGGGTAATGTATTTGGGTTCAAAGCG
CTGCGCGCTCTACGTCTGGAAGATCTGCGAATCCCTACTGCGTATGTTAAAACTTTCCAAGGACCGCCTCATGGTATCCAAGTTGAAAGAGA
TAAATTGAACAAGTATGGTCGTCCCCTGTTGGGATGTACTATTAAACCTAAATTGGGGTTATCTGCTAAAAACTACGGTAGAGCAGTTTATG
AATG
```

Requires input taxonomy file containing lineages associated with sequences, typically in Metaxa2-compatible format. For certain classifiers, such as Metaxa2, higher resolution ranks can be omitted for cases of ambiguous open nomenclature. The curation toolkit is made to accommodate this but such entries may need to be removed for alternative classifiers such as SINTAX or RDP.

```
NC_035751.1	k__Eukaryota;p__Streptophyta;c__Liliopsida;o__Poales;f__Poaceae;g__Miscanthus;s__Miscanthus junceus;
EU697437.1	k__Eukaryota;p__Streptophyta;c__Jungermanniopsida;o__Metzgeriales;f__Aneuraceae;g__Aneura;
KF284759.1	k__Eukaryota;p__Streptophyta;c__Liliopsida;o__Alismatales;f__Araceae;g__Colocasia;
```

Optionally, input taxonomies can come directly from the Taxonomizr R module, such as in the example shown below, if ‘-tf True’ is declared

```
"  13597","L11686.1","13597","Eukaryota","Streptophyta",NA,"Lamiales","Oleaceae","Ligustrum","Ligustrum vulgare"
"  13525","L11684.1","13525","Eukaryota","Streptophyta",NA,"Gentianales","Gentianaceae","Exacum","Exacum affine"
```
