## Supplemental Data for "MetaCurator: A hidden Markov model-based toolkit for extracting and curating sequences from taxonomically-informative genetic markers"

**Supplemental Table S1:** Universal primer sets of which these databases are representative.

| Marker | Primers | Source |
|---|---|---|
| *rbcL* | rbcL2: TGGCAGCATTYCGAGTAACTC<br>rbcLa-R: GTAAAATCAAGTCCACCRCG | Palmieri et al., 2009<br>Kress and Erickson 2007 |
| *trnL* | A49325: CGAAATCGGTAGACGCTACG<br>B49863: GGGGATAGAGGGACTTGAAC | Taberlet et al., 1991 |
| ITS2 | ITS-S2F: ATGCGATACTTGGTGTGAAT<br>ITS4R: TCCTCCGCTTATTGATATGC | White et al. 1990<br>Chen et al. 2010 |
| COI | BF2: GCHCCHGAYATRGCHTTYCC<br>BR2: TCDGGRTGNCCRAARAAYCA | Elbrecht and Leese 2017 |

**Supplemental Table S2:** Proportion of taxa retained during database curation for sequences belonging to each respective marker according to NCBI annotations. Since many markers represent long sequence regions which contain multiple commonly used barcodes with varying levels of overlap, some loss of taxa is inevitable. For example, researchers often use two very short *trnL* barcodes, which would not have been extracted in this study since we extracted one of the longest *trnL* barcodes and required a percent profile match coverage threshold of 85 percent of the length of the marker.

| Study System | Marker | Orders | Families | Genera | Species |
|---|---|---|---|---|---|
| Plant | *trnL* | 0.664 | 0.658 | 0.756 | 0.600 |
| | *rbcL* | 1.000 | 0.978 | 0.929 | 0.753 |
| | *trnH* | 0.907 | 0.818 | 0.686 | 0.522 |
| | ITS2 | 0.926 | 0.860 | 0.704 | 0.513 |
| Arthropod | COI | 0.991 | 0.945 | 0.877 | 0.852 |

**Supplemental Table S3:** Percent increase in database richness between this study and Richardson et al. (2018 and 2019).

| | Orders | Families | Genera | Species |
|---|---|---|---|---|
| *trnL* | 27.4 | 21.4 | 42.7 | 180.9 |
| *rbcL* | 12.2 | 11.0 | 46.3 | 193.8 |
| COI | 6.5 | 32.1 | 45.2 | 69.4 |

**Supplemental Figure S1:** Sequences yielded per iteration per round for each database curation. Panel labels, 1 through 5, represent extraction rounds which varied in in percent profile HMM length coverage required, from 100 to 85 percent. In general, high-conservation markers, like *rbcL*, require few iterations per round as most available sequences are extracted in the first round. Nonetheless, we recommend multiple iterations per round even when using high-conservation markers since the few sequences which will be added may represent a significant proportion of the phylogenetic diversity of the resulting database.

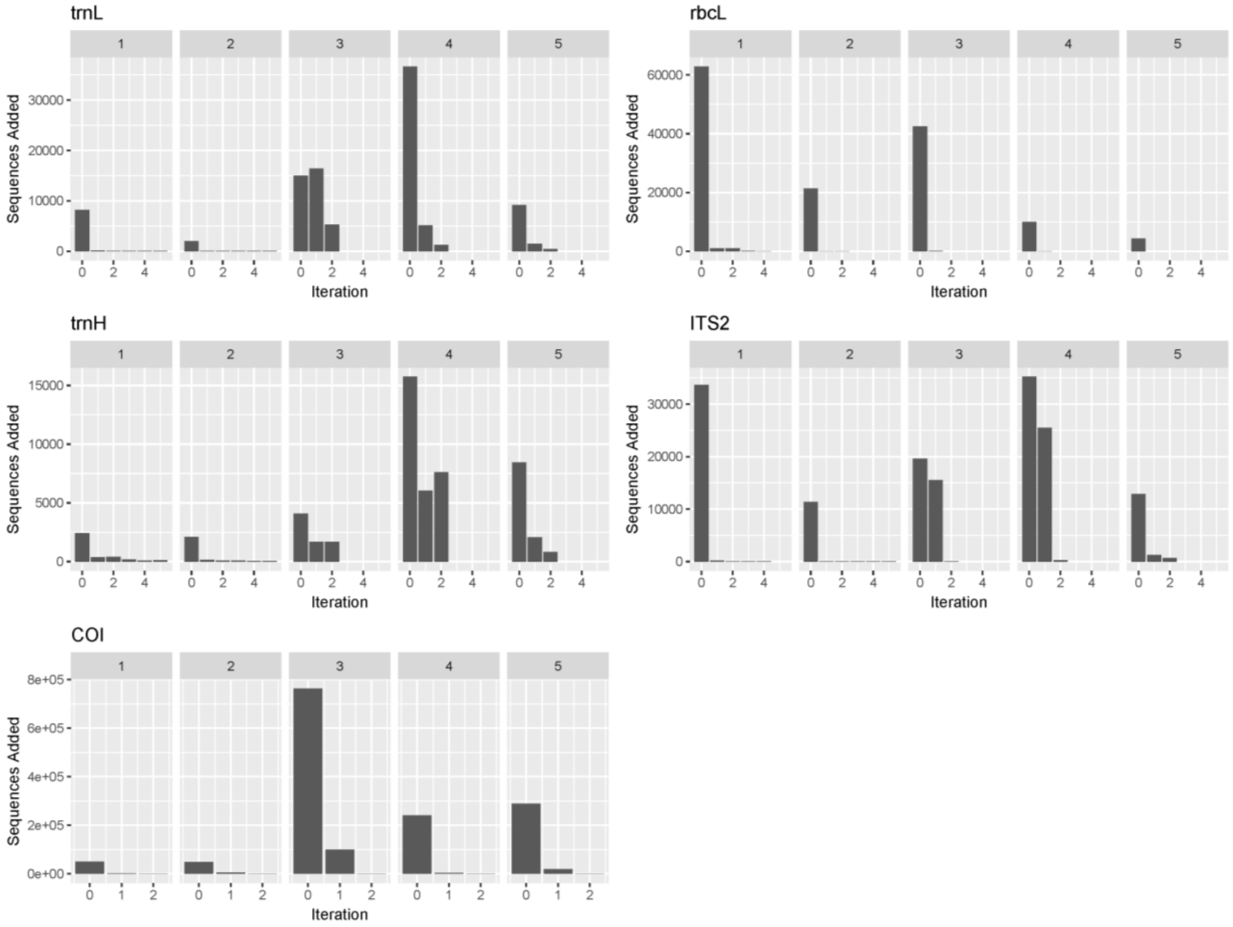

**Supplemental Figure S2**: Multiple sequence alignment of 50 sequences randomly drawn from the *rbcL* database produced with Muscle and annotated with BoxShade (v3.21).

```
AF419078.1   1 TGGCAGCATTCCGAATGACTCCGCAACCCGGCGTGCCTGCTGAGGAAGCGGGAGCCGCGG
L34816.1     1 --GCAGCTTTCCGTATGACTCCTCAGGCCTGTGTACCACCTGAAGAAGCTGGTGCAGCTG
KP686040.1   1 TGGCAGCTTTTCGAATGACTCCGCAACCTGGAGTTCCACCAGAAGAATGTGGCGCTGCTG
KJ446802.1   1 --GCAGCATTCCGTATGACTCCTCAACCAGGTGTTCCACCAGAAGAAGCTGGTGCAGCGG
FN432103.1   1 TGGCTGCATTCCGTATGACTCCTCAACCTGGAGTTCCTCCAGAAGAAGCTGGAGCAGCTG
KX656068.1   1 --GCAGCCTTCCGAATGACCCCACAACCCGGAGTACCAGCTGAAGAAGCCGGAGCTGCGG
JN572331.1   1 --GCAGCCTTCCGGATGACCCCACAACCCGGAGTACCAGCCGAGGAGGCTGGAGCTGCGG
U05608.1     1 TGGCAGCGTTCCGAATGACTCCGCAACCTGGAGTACCACCAGAGGAAGCTGGAGCTGCAG
KY089068.1   1 --GCAGCATTTCGAATGACTCCTCAACCAGGCGTACCTGCTGAAGAGGCAGGGGCTGCAG
AY452692.1   1 --GCAGCATTTCGAATGACTCCTCAACCAGGAGTGCCAGCTGAAGAAGCAGGAGCTGCAG
JF940531.1   1 TGGCGGCATTCCGAGTAACTCCTCAACCTGGGGTGCCGCCCGAGGAAGCGGGAGCAGCAG
GQ408938.1   1 --GCCGCATTCCGAGTAACTCCTCAACCTGGGGTTCCCCCTGAGGAAGCGGGCGCTGCAG
HQ901564.1   1 TGGCAGCATTCCGAGTAACCCCGCAACCCGGAGTTCCACCTGAGGAAGCAGGGGCCGCAG
LN907888.1   1 TGGCAGCATTCCGAGTAACTCCTCAGCCCGGGGTTCCGCCTGAAGAAGCAGGGGCTGCAG
KR736525.1   1 -GGCAGCATTCCGAGTAACTCCTCAGCCCGGGGTTCCGCCTGAAGAAGCAGGGGCTGCAG
KY627310.1   1 -GGCAGCATTCCGAGTAACTCCTCAGCCCGGGGTTCCGCCTGAAGAAGCAGGGGCTGCAG
KC483293.1   1 -GGCTGCATTCCGAGTTACTCCTCAACCTGGAGTTCCAGCTGAGGAAGCAGGGGCCGCGG
KJ204348.1   1 TGGCAGCATTCCGAGTAACTCCTCAACCCGGAGTTCCACCTGAGGAAGCGGGGGCCGCGG
KT458050.1   1 TGGCAGCATTCAGAGTAACTCCTCAACCTGGAGTTCCGCCTGAGGAAGCAGGGGCTGCAG
M77031.1     1 TGGCAGCATTCCGAGTAACTCCTCAACCTGGAGTTCCGCCTGAGGAAGCAGGAGCTGCGG
AY875228.1   1 TGGCAGCATTCCGAGTAACTCCTCAACCTGGAGTTCCGTCAGAAGAAGCAGGAGCCGCAG
DQ907755.1   1 TGGCGGCATTCCGAGTAACTCCTCAACCCGGAGTTCCAGCCGAAGAAGCAGGTGCCGCAG
JQ933274.1   1 TGGCAGCATTCCGAGTAAGTCCTCAACCCGGAGTTCCACCCGAAGAAGCAGGGGCCGCAG
EF178544.1   1 TGGCAGCGTTCCGAGTAACTCCTCAACCCGGAGTCCCCCCTGAAGAAGCAGGAGCTGCAG
AB586281.1   1 TGGCAGCGTTCCGAGTAACTCCTCAACCTGGAGTCCCTCCTGAAGAAGCAGGAGCTGCAG
AM999843.1   1 TGGCAGCGTTCCGAGTAACTCCTCAACCCGGAGTCCCCCCTGAAGAAGCAGGAGCTGCAG
AB872559.1   1 TGGCAGCATTCCGAGTAACTCCTCAACCCGGGGTTCCGCCCGAGGAAGCCGGGGCCGCGG
EU643727.1   1 TGGCAGCCTTTCGAGTCACTCCTCAACCCGGAGTTCCCCCGGAAGAAGCCGGAGCCGCAG
AY298820.1   1 TGGCAGCATTCCGAGTAACTCCTCAACCTGGAGTTCCGCCGGAAGAAGCAGGGGCTGCGG
HM640507.1   1 TGGCAGCATTCCGAGTAACTCCTCAACCTGGAGTTCCCGCTGAAGAAGCAGGGGCTGCGG
KC704911.1   1 TGGCAGCATTCCGAGTAACTCCTCAACCGGGAGTTCCGCCTGAAGAAGCGGGGGCTGCGG
KU748062.1   1 -GGCAGCATTCCGAGTAACCCCTCAACCGGGAGTTCCACCTGAAGAAGCGGGGGCTGCGG
DQ875838.1   1 TGGCAGCATTCCGGGTAACTCCTCAACCTGGAGTTCCTCCTGAGGAAGCGGGAGCAGCGG
EU840324.1   1 TGGCAGCATTTCGAGTAACTCCTCAACCTGGAGTTCCACCAGAAGAAGCAGGGGCCGCGG
JX571893.1   1 TGGCAGCATTTCGAGTAACTCCTCAACCAGGAGTTCCACCGGAAGAAGCAGGGGCCGCGG
AB586309.1   1 TGGCAGCATTCCGAGTAACTCCTCAACCTGGAGTTCCGCCTGAAGAAGCAGGTGCCGCGG
KX397733.1   1 -GGCAGCATTCCGAGTAACTCCTCAACCTGGAGTTCCGCCTGAAGAAGCAGGTGCCGCGG
JF944669.1   1 TGGCAGCATTCCGAGTAACTCCTCAACCGGGAGTTCCACCCGAGGAAGCAGGGGCCGCTG
DQ535789.1   1 TGGCAGCATTCCGAGTAACTCCTCAACCTGGAGTTCCACCCGAGGAAGCAGGGGCTGCTG
KM606572.1   1 TGGCAGCATTCCGAGTAACTCCTCAACCGGGAGTTCCACCTGAAGAAGCAGGGGCCGCGG
HQ260780.1   1 --GCAGCATTCCGAGTAACTCCTCAACCTGGAGTTCCGCCGGAAGAAGCGGGGGCTGCGG
MF065382.1   1 --------------------------------------------GAAGCAGGGGCCGCAG
AY570449.1   1 TGGCAGCATTCCGAGTAACTCCTCAACCCGGAGTTCCGCCTGAAGAAGCAGGGGCCGCGG
AF190445.1   1 TGGCAGCATTCCGAGTAACTCCCCAACCTGGAGTTCCACCGGAAGAAGCAGGGGCCGCGG
HM849885.1   1 --GCAGCATTTCGAGTAACTCCTCAACCTGGAGTTCCACCTGAAGAAGCAGGGGCCGCAG
MG221719.1   1 -GGCAGCATTTCGAGTAACTCCTCAACCTGGAGTTCCGCCTGAAGAAGCAGGGGCCGCAG
JQ933508.1   1 TGGCAGCATTCCGAGTAACTCCTCAACCTGGAGTTCCACCTGAAGAAGCGGGGGCCGCGG
JF308648.1   1 TGGCAGCATTCCGAGTAACTCCTCAACCTGGAGTTCCACCCGAGGAAGCAGGGGCCGCAG
AB744227.1   1 TGGCAGCATTCCGAGTAACTCCTCAACCTGGAGTTCCACCTGAAGAAGCAGGGGCCGCGG
AM117226.1   1 TGGCAGCATTCCGAGTAACTCCTCAACCGGGAGTTCCACCTGAAGAAGCAGGGGCCGCGG
```

| | | |
|---|---|---|
| AF419078.1 | 61 | TAGCCGCGGAGTCCTCCACCGGCACGTGGACCACCGTCTGGACTGACGGGCTTACCAATC |
| L34816.1 | 59 | TAGCAGCTGAATCATCAACAGGTACTTGGACTACAGTATGGACTGATGGTCTAACTAGCC |
| KP686040.1 | 61 | TTGCTGCTGAATCTTCTACTGGTACATGGACTACAGTTTGGACGGATGGTTTAACAAGCT |
| KJ446802.1 | 59 | TAGCAGCAGAATCATCAACAGGTACTTGGACAACTGTATGGACAGACGGTTTAACGAGTT |
| FN432103.1 | 61 | TTGCAGCAGAGTCTTCCACCGGAACATGGACTACTGTTTGGACTGATGGTCTTACTAGCC |
| KX656068.1 | 59 | TAGCTGCGGAATCCTCCACGGGTACGTGGACCACTGTATGGACAGATGGATTGACCAATC |
| JN572331.1 | 59 | TAGCTGCGGAATCCTCAACGGGTACGTGGACCACTGTATGGACAGACGGGTTGACCAGCC |
| U05608.1 | 61 | TAGCTGCGGAATCTTCCACAGGTACATGGACCACAGTATGGACAGATGGGCTTACTAGCC |
| KY089068.1 | 59 | TAGCTGCGGAATCTTCCACTGGTACATGGACCACTGTTTGGACTGATGGACTTACCAGTC |
| AY452692.1 | 59 | TAGCTGCAGAATCTTCTACTGGTACATGGACTACTGTTTGGACTGATGGCCTTACTAGTC |
| JF940531.1 | 61 | TAGCTGCTGAATCTTCCACCGGTACATGGACCACTGTTTGGACCGATGGACTTACCAGTC |
| GQ408938.1 | 59 | TAGCTGCCGAATCTTCTACGGGTACATGGACAACGGTTTGGACTGATGGACTTACCAGCC |
| HQ901564.1 | 61 | TAGCTGCCGAATCTTCTACTGGTACATGGACAACCGTGTGGACTGATGGACTTACTAGTC |
| LN907888.1 | 61 | TAGCTGCGGAATCTTCTACTGGTACATGGACAACTGTTTGGACTGATGGACTTACCAGTC |
| KR736525.1 | 60 | TAGCTGCGGAATCTTCTACTGGTACATGGACAACTGTTTGGACTGATGGACTTACCAGTC |
| KY627310.1 | 60 | TAGCTGCGGAATCTTCTACTGGTACATGGACAACTGTTTGGACTGATGGACTTACCAGTC |
| KC483293.1 | 60 | TAGCTGCCGAATCTTCTACTGGTACATGGACAACTGTGTGGACCGATGGACTTACCAGCC |
| KJ204348.1 | 61 | TAGCTGCTGAATCTTCTACCGGTACATGGACAACCGTGTGGACCGATGGGCTTACTAGTC |
| KT458050.1 | 61 | TAGCTGCCGAATCTTCTACTGGTACCTGGACAACTGTGTGGACCGATGGGCTTACCAGCC |
| M77031.1 | 61 | TGGCTGCCGAATCTTCCACTGGTACATGGACAACTGTGTGGACCGATGGACTTACCAGCC |
| AY875228.1 | 61 | TAGCTGCCGAATCTTCTACTGGTACATGGACAACTGTATGGACCGACGGACTTACCAGTC |
| DQ907755.1 | 61 | TAGCTGCGGAATCTTCTACTGGTACATGGACAACTGTATGGACCGACGGACTTACCAGTC |
| JQ933274.1 | 61 | TAGCCGCCGAATCTTCGACTGGTACATGGACAACTGTATGGACCGACGGACTTACCAATC |
| EF178544.1 | 61 | TAGCTGCCGAATCTTCTACTGGTACATGGACAACTGTTTGGACTGATGGACTTACCAGTC |
| AB586281.1 | 61 | TAGCGGCGGAATCTTCTACTGGTACATGGACAACTGTTTGGACTGATGGACTTACCAGTC |
| AM999843.1 | 61 | TAGCGGCGGAATCTTCTACTGGTACATGGACAACTGTTTGGACTGATGGACTTACCAGTC |
| AB872559.1 | 61 | TAGCTGCGGAATCTTCTACTGGTACATGGACAACTGTGTGGACCGATGGGCTTACCAGCC |
| EU643727.1 | 61 | TAGCTGCCGAATCGTCTACTGGTACATGGACAACTGTGTGGACCGATGGGCTTACGAGCC |
| AY298820.1 | 61 | TAGCAGCCGAGTCTTCTACTGGTACATGGACAACCGTTTGGACTGATGGACTTACCAGTC |
| HM640507.1 | 61 | TAGCTGCCGAATCTTCTACTGGTACATGGACAACTGTGTGGACCGATGGACTTACCAGTC |
| KC704911.1 | 61 | TAGCTGCCGAATCTTCTACTGGTACATGGACAACTGTGTGGACTGATGGACTTACCAGTC |
| KU748062.1 | 60 | TAGCTGCCGAATCTTCTACTGGTACATGGACAACTGTGTGGACTGATGGACTTACCAGTC |
| DQ875838.1 | 61 | TAGCTGCCGAATCTTCTACTGGTACATGGACAACTGTGTGGACCGATGGACTTACCAGCC |
| EU840324.1 | 61 | TAGCTGCCGAATCTTCTACTGGTACATGGACAACTGTGTGGACCGATGGACTTACCAGCC |
| JX571893.1 | 61 | TAGCTGCCGAATCTTCTACTGGTACATGGACAACTGTGTGGACCGATGGACTTACCAGCC |
| AB586309.1 | 61 | TAGCTGCTGAATCTTCTACTGGTACATGGACAACTGTGTGGACCGATGGGCTTACCAGTC |
| KX397733.1 | 60 | TAGCTGCTGAATCTTCTACTGGTACATGGACAACTGTGTGGACCGATGGGCTTACCAGTC |
| JF944669.1 | 61 | TAGCTGCTGAATCTTCTACTGGTACATGGACAACTGTGTGGACCGATGGACTTACCAGTC |
| DQ535789.1 | 61 | TAGCTGCTGAATCTTCTACTGGTACATGGACAACTGTGTGGACCGATGGGCTTACCAGTC |
| KM606572.1 | 61 | TAGCTGCGGAATCTTCTACTGGTACATGGACAACAGTGTGGACTGATGGACTTACTAGCC |
| HQ260780.1 | 59 | TAGCTGCCGAATCTTCTACAGGTACATGGACAACTGTGTGGACCGATGGACTTACCAGTC |
| MF065382.1 | 17 | TAGCTGCCGAATCTTCTACTGGTACATGGACAACCGTGTGGACCGATGGACTTACCAGCC |
| AY570449.1 | 61 | TAGCTGCCGAATCTTCTACTGGTACATGGACAACTGTGTGGACCGATGGACTTACCAGCC |
| AF190445.1 | 61 | TAGCTGCCGAGTCTTCTACTGGTACATGGACAACTGTATGGACGGATGGACTTACCAGTC |
| HM849885.1 | 59 | TAGCTGCCGAATCTTCTACTGGTACATGGACAACTGTATGGACCGATGGACTTACGAGCC |
| MG221719.1 | 60 | TAGCTGCCGAATCTTCTACTGGTACATGGACAACTGTGTGGACCGATGGACTTACGAGCC |
| JQ933508.1 | 61 | TAGCTGCCGAATCTTCTACTGGTACATGGACCACTGTGTGGACCGATGGACTTACCAGCC |
| JF308648.1 | 61 | TAGCTGCCGAATCTTCTACTGGTACATGGACAACTGTGTGGACCGATGGACTTACCAGCC |
| AB744227.1 | 61 | TAGCTGCCGAATCTTCTACTGGTACATGGACAACTGTATGGACCGATGGACTTACCAGTC |
| AM117226.1 | 61 | TAGCTGCCGAATCTTCTACTGGTACATGGACAACTGTGTGGACCGATGGGCTTACCAGCC |

```
AF419078.1  121 TTGATCGTTATAAGGGTCGATGCTATGGCATTGAAAAAGTGGCAGGGGAAAAGGATCAAT
L34816.1    119 TTGATCGTTATAAGGGTCGTTGTTACGATATTGAACCAGTTCCGGGTGAAGATGAGCAGT
KP686040.1  121 TAGATAGTTATAAAGGACGTTGTTATGATTTAGAACCGGTGAAAGGGGAAGAAAATCAAT
KJ446802.1  119 TAGATCGTTACAAAGGTCGTTGTTATGACATCGAGCCAGTTCCCGGTGAAGAAAACCAAT
FN432103.1  121 TAGATCGTTACAAAGGTCGTTGTTACGATATCGAACCTGTACCTGGAGAAGAAAACCAGT
KX656068.1  119 TTGACCGTTACAAGGGCCGATGCTACGACATTGAACCCGTCGCTGGGGAAGAAAACCAGT
JN572331.1  119 TTGACCGTTACAAGGGTCGATGCTATGATATTGAACCTGTCGCTGGAGAAGAAAACCAGT
U05608.1    121 TTGATCGTTATAAGGGTCGATGCTACGATATCGAACCTGTTCCTGGAAGCGATAATCAAT
KY089068.1  119 TTGATCGTTATAAAGGACGATGCTATGATCTTGAAGCAGTTCCTGGAGAAGAGAATCAAT
AY452692.1  119 TTGATCGTTATAAAGGACGATGCTATGATCTTGAAGCAGTTGCTGGAGAAGATAATCAAT
JF940531.1  121 TTGATCGTTACAAAGGGCGATGCTATGACATCGAGCCCGTTGCTGGAGAGGAAAGTCAAT
GQ408938.1  119 TTGATCGTTACAAAGGACGATGCTATCACATCGAGCCCGTTATTGGAGAGGAAAATCAAT
HQ901564.1  121 TAGATCGTTACAAAGGACGATGCTACCACATCGAGCCTGTTATTGGGGAGGAAAATCAAT
LN907888.1  121 TTGATCGTTACAAAGGACGATGCTATCACATCGAGCCCGTTCCTGGGGATGAAAATCAAT
KR736525.1  120 TTGATCGTTACAAAGGACGATGCTATCACATCGAGCCCGTTCCTGGGGAGGCAGATCAAT
KY627310.1  120 TTGATCGTTACAAAGGACGATGCTATCACATCGAGCCCGTTCCTGGGGACGAAGAGCAAT
KC483293.1  120 TTGATCGTTACAAAGGAAGATGCTACCACATCGAGCCCGTTGCTGGAGAAGACAATCAAT
KJ204348.1  121 TGGATCGTTACAAAGGACGCTGCTATCACATCGAGCCCGTTGCTGGAGAAGAAAATCAAT
KT458050.1  121 TTGATCGTTATAAAGGAAGATGCTACCACATCGAGCCTGTTGCGGGAGAAGAGAATCAAT
M77031.1    121 TCGATCGTTACAAAGGACGATGCTACCACATCGAGCCAGTTGCTGGGGAGGAAAATCAAT
AY875228.1  121 TTGATCGTTACAAAGKACGATGCTACCACATCGATGCCGTTCCTGGAGAAGACAATCAAT
DQ907755.1  121 TTGATCGTTACAAAGGCCGATGCTACCACATCGAGCCTGTTGCTGGAGACGAAAATCAAT
JQ933274.1  121 TTGATCGTTACAAAGGACGATGCTACCACATCGAGCCCGTTGCTGGAGAAGAAAATCAAT
EF178544.1  121 TTGATCGTTACAAAGGACGATGCTACCACATCGAGCCTGTTGTTGGAGAAGAAAATCAAT
AB586281.1  121 TTGATCGTTACAAAGGGCGATGCTATCATATCGAACCTGTTGCTGGAGAAGAAAATCAAT
AM999843.1  121 TTGATCGTTACAAAGGGCGATGCTATCATATCGAGCCTGTTGTTGGAGAAGAAAATCAAT
AB872559.1  121 TTGATCGTTATAAAGGACGATGCTACAACATTGAGCCCGTTGCTGGAGAAGAAAATCAAT
EU643727.1  121 TTGATCGTTACAAAGGGAGATGCTATCACATTGAGCCCGTTGCCGGAGAAGAAAATCAAT
AY298820.1  121 TTGATCGTTACAAAGGAAGATGCTACCACATCGAGCCCGTTGTCGGAGAGGAAAATCAAT
HM640507.1  121 TTGATCGTTACAAAGGACGATGCTACCACATTGAGGCCGTTGTTGGAGAAGAAAATCAAT
KC704911.1  121 TTGATCGTTACAAAGGACGATGCTACCACATCGAGCCCGTTGCTGGGGAGGAAAATCAAT
KU748062.1  120 TTGATCGTTACAAAGGACGATGCTACCACATCGAGGCCGTTGTTGGGGAGGAAAATCAAT
DQ875838.1  121 TTGATCGTTACAAAGGACGATGCTACCACATTGAGCCGGTTGCTGGAGAAGAAAATCAAT
EU840324.1  121 TTGATCGTTACAAAGGACGATGCTACCACATCGAGCCTGTTCCTGGAGAAGAAAGTCAGT
JX571893.1  121 TTGATCGTTACAAAGGACGATGCTACCACATTGAGCCTGTTACTGGAGAAGAAAATCAAT
AB586309.1  121 TTGATCGTTACAAAGGACGATGCTACCACATCGAGCCCGTTGCTGGAGAAGAAAATCAAT
KX397733.1  120 TTGATCGTTACAAAGGACGATGCTACCACATCGAGCCCGTTCCTGGAGAAGAAAGTCAAT
JF944669.1  121 TTGATCGTTACAAAGGACGATGCTATGACATCGAGCCTGTTCCTGGAGAAGAAAGTCAAT
DQ535789.1  121 TTGATCGTTACAAAGGACGATGCTACGACATCGAGCCTGTTGCTGGAGAAGAAAATCAAT
KM606572.1  121 TTGATCGTTACAAAGGGCGATGCTACCACATCGAGCCTGTTGCTGGAGACGAAAATCAAT
HQ260780.1  119 TTGATCGTTACAAAGGACGATGCTACAACATTGAGCCTGTTGCTGGGGAAGAAAATCAAT
MF065382.1   77 TTGATCGTTACAAAGGGCGATGCTACCACATCGAGCCCGTTCCTGGAGAAGCAGATCAAT
AY570449.1  121 TTGATCGTTACAAAGGGCGATGCTACCACATTGAGCCCGTTCCTGGAGAAAAAGATCAAT
AF190445.1  121 TTGATCGTTACAAAGGGCGATGCTACCACATCGAGCCAGTTCCTGGAGAAGAAGATCAAT
HM849885.1  119 TTGATCGTTACAAAGGGCGATGCTATGGAATTGAGCCTGTTCCTGGAGAAGAGAATCAAT
MG221719.1  120 TTGATCGTTACAAAGGGCGATGCTATGGAATCGAGCCTGTTCCTGGAGAAGAAAATCAAT
JQ933508.1  121 TTGATCGTTACAAAGGGCGCTGCTACGGAATCGAGCCCGTTGCTGGAGAAGAAAATCAAT
JF308648.1  121 TTGATCGTTACAAAGGGCGATGCTACAACATCGAGCCCGTTGCTGGAGAAGAAAATCAAT
AB744227.1  121 TTGATCGTTACAAAGGGCGATGCTACCGCATCGAGCGTGTTCTTGGAGAAAAAGATCAAT
AM117226.1  121 TTGATCGTTACAAAGGGCGATGCTACCACATCGAGCCAGTTCCTGGAGAAGAAGATCAAT
```

```
AF419078.1  181 ATATTGCCTACGTGGCCTATCCCTTGGATCTGTTCGAGGAAGGCTCCGTTACCAATATGC
L34816.1    179 ACATTTGTTACATTGCTTACCCTATTGATCTTTTTGAAGAAGGTTCTGTAACTAACCTGT
KP686040.1  181 ATATAGCTTATGTGGCTTATCCTCTTGATTTATTTGAGGAAGGATCGGTTACTAATTTAT
KJ446802.1  179 ACATTGCATATATTGCATACCCTTTAGACCTTTTTGAAGAAGGATCTGTAACCAATTTAT
FN432103.1  181 ACATCGCATACGTAGCTTATCCTCTAGATCTATTCGAAGAAGGTTCCGTCACCAACCTTT
KX656068.1  179 ATATCGCGTATGTAGCTTATCCTTTGGATCTATTCGAAGAAGGTTCTGTCACCAATTTGT
JN572331.1  179 ATATCGCATATGTAGCTTATCCTTTGGATCTATTCGAAGAAGGTTCTGTCACCAACTTGT
U05608.1    181 ATATTGCATATGTAGCTTATCCTTTAGATCTCTTTGAAGAAGGATCTGTTACGAATCTAT
KY089068.1  179 ATATTGCTTATGTTGCTTACCCATTAGATCTATTTGAAGAAGGTTCTGTTACCAATTTAT
AY452692.1  179 ATATTGCTTATGTTGCTTACCCATTAGATTTATTTGAAGAAGGTTCTGTTACCAATTTAT
JF940531.1  181 TTATTGCCTATGTAGCTTACCCCTTAGACCTTTTCGAAGAAGGTTCTGTTACTAACTTGT
GQ408938.1  179 ATATTGCCTATGTAGCTTATCCTTTAGACCTTTTTGAAGAGGGTTCTGTTACTAACATGT
HQ901564.1  181 ATTTTTGTTATGTAGCTTATCCTTTGGACCTTTTTGAAGAGGGTTCCGTTACCAACATGT
LN907888.1  181 ATATCTGTTATGTAGCTTACCCATTAGACCTATTTGAAGAGGGTTCTGTTACTAACATGT
KR736525.1  180 ATATCTGTTATATAGCTTATCCATTAGACCTATTTGAAGAGGGTTCTGTTACTAACATGT
KY627310.1  180 ATATCTGTTATGTAGCTTATCCATTAGACCTATTTGAAGAGGGTTCTGTTACTAACATGT
KC483293.1  180 ATATTTGTTATGTAGCTTATCCTTTAGACCTTTTTGAAGAAGGTTCTGTTACTAACATGT
KJ204348.1  181 TTATTGCTTATGTAGCTTACCCTTTAGACCTTTTTGAAGAAGGTTCCGTTACTAATATGT
KT458050.1  181 ATATATGTTATGTAGCTTACCCCTTAGACCTTTTTGAAGAGGGTTCTGTTACTAATATGT
M77031.1    181 ATATTGCTTATGTAGCTTACCCTTTGGATCTTTTTGAAGAAGGTTCTGTTACTAACATGT
AY875228.1  181 ATATYTGTTATGTAGCTTACCCCTTAGCACCTYTTGAAGAAGGTTCTGTTACTAATATGT
DQ907755.1  181 ATATTTGTTATGTAGCTTACCCCTTAGACCTTTTTGAGGAAGGCTCCGTTACTAACATGT
JQ933274.1  181 ATATTTGTTATGTAGCTTACCCCTTAGATCTTTTTGAGGAAGGCTCTGTTACTAACATGT
EF178544.1  181 ATATTGCCTATGTAGCTTATCCTTTAGACCTTTTCGAAGAGGGTTCTGTTACTAACATGT
AB586281.1  181 ATATTGCCTATATAGCTTATCCTTTAGACCTTTTCGAAGAAGGTTCTGTTACTAACATGT
AM999843.1  181 ATATTGCCTATGTAGCTTATCCTTTAGACCTTTTTGAAGAAGGTTCTGTTACTAACATGT
AB872559.1  181 ATATATGTTATGTAGCTTACCCTTTAGACCTTTTTGAAGAGGGTTCTGTTACTAACATGT
EU643727.1  181 TTATTGCTTATGTAGCTTACCCATTAGACCTTTTTGAAGAAGGTTCTGTTACTAACATGT
AY298820.1  181 ATATTGCTTATGTAGCTTATCCTTTAGACCTTTTTGAAGAAGGTTCTGTTACTAACATGT
HM640507.1  181 ATATTGCTTATGTAGCTTATCCTTTAGACCTTTTTGAAGAAGGTTCTGTTACTAACATGT
KC704911.1  181 ATATTGCTTATGTAGCTTATCCTTTAGATCTTTTTGAAGAAGGTTCTGTTACTAACATGT
KU748062.1  180 ATATTGCTTATGTAGCTTATCCATTAGACCTTTTTGAAGAAGGTTCTGTTACTAACATGT
DQ875838.1  181 ATATTGCTTATGTAGCTTACCCTTTAGACCTTTTTGAAGAAGGTTCTGTTACTAACATGT
EU840324.1  181 TTATTGCTTATGTAGCTTACCCATTAGACCTTTTTGAAGAAGGTTCTGTTACTAACATGT
JX571893.1  181 ATATTTGTTATGTAGCTTACCCTTTAGATCTTTTTGAAGAAGGTTCTGTTACTAACATGT
AB586309.1  181 ATATTGCTTATGTAGCTTATCCCTTAGACCTTTTTGAAGAAGGTTCTGTTACTAACATGT
KX397733.1  180 TTATTGCTTATGTAGCTTATCCCTTAGACCTTTTTGAAGAAGGTTCTGTTACTAACATGT
JF944669.1  181 TTATTGCTTATGTAGCTTATCCCCTAGACCTTTTTGAAGAAGGTTCTGTTACTAACATGT
DQ535789.1  181 ATATTGCTTATGTAGCTTATCCCTTAGACCTTTTTGAAGAAGGTTCTGTTACTAACATGT
KM606572.1  181 ATATTGCTTATGTAGCTTATCCTTTAGACCTTTTTGAAGAAGGTTCTGTTACTAACATGT
HQ260780.1  179 ATATTTGTTATGTAGCTTACCCCTTAGACCTTTTTGAAGAAGGGTCTGTTACTAATATGT
MF065382.1  137 ATATCTGTTATGTAGCTTACCCTTTAGACCTTTTTGAAGAAGGTTCTGTTACTAACATGT
AY570449.1  181 ATATCTGTTATGTAGCTTACCCTTTAGACCTTTTTGAAGAAGGTTCTGTTACTAACATGT
AF190445.1  181 TTATTGCTTATGTAGCTTACCCATTAGACCTTTTTGAAGAAGGTTCTGTTACTAACATGT
HM849885.1  179 ATATTGCTTATGTAGCTTACCCATTAGACCTTTTTGAAGAAGGTTCTGTTACTAACATGT
MG221719.1  180 TTATTGCTTATGTAGCTTACCCATTAGACCTTTTTGAAGAAGGTTCTGTTACTAACATGT
JQ933508.1  181 ATATCGCTTATGTAGCTTACCCATTAGACCTTTTTGAAGAAGGTTCTGTTACTAACATGT
JF308648.1  181 ATATTGCTTATGTAGCTTACCCCTTAGACCTTTTTGAAGAAGGTTCTGTTACTAACATGT
AB744227.1  181 ATATTGCTTATGTAGCTTACCCTTTAGACCTTTTTGAAGAAGGTTCCGTTACCAATATGT
AM117226.1  181 ATATTGCTTATGTAGCTTACCCCTTAGACCTTTTTGAAGAAGGTTCTGTTACTAACATGT
```

```
AF419078.1  241 TCACCTCCATAGTAGGTAACGTTTTCGGGTTTAAGGCCCTGCGGNCTCTGCGCCTGGAAG
L34816.1    239 TCACATCAATCGTAGGTAACGTATTTGGTTTCAAAGCTCTTCGTGCTCTTCGTCTTGAAG
KP686040.1  241 TTACATCAATTGTTGGAAATGTGTTTGGTTTTAAAGCATTAAGAGCTCTTCGATTAGAAG
KJ446802.1  239 TTACTTCAATTGTAGGTAACGTTTTTGGTTTCAAAGCTCTTCGTGCTTTACGTTTAGAAG
FN432103.1  241 TTACTTCCATTGTAGGTAACGTTTTTGGATTTAAAGCTCTTAGAGCTCTACGTCTAGAAG
KX656068.1  239 TCACCTCCATTGTAGGTAATGTTTTTGGATTTAAGGCTCTACGCGCCTTACGCTTGGAAG
JN572331.1  239 TCACCTCCATCGTAGGTAATGTTTTTGGATTTAAGGCTCTACGCGCTCTACGCTTGGAAG
U05608.1    241 TCACTTCGATTGTAGGTAACGTTTTTGGATTTAAAGCCTTACGTGCCCTACGGTTGGAAG
KY089068.1  239 TTACTTCTATTGTTGGTAATGTTTTCGGATTTAAAGCTTTACGAGCTTTACGTCTAGAAG
AY452692.1  239 TTACTTCTATTGTTGGTAATGTTTTTGGATTTAAAGCTTTACGAGCTTTACGTCTAGAAG
JF940531.1  241 TCACTTCCATTGTAGGTAATGTATTTGGATTCAAGGCCCTACGGGCTTTACGTTTGGAAG
GQ408938.1  239 TTACTTCGATTGTGGGCAATGTATTTGGTTTCAAAGCGCTACGAGCTCTACGCCTGGAGG
HQ901564.1  241 TTACTTCCATTGTAGGTAATGTATTTGGGTTTAAAGCTCTACGAGCTCTACGTTTGGAGG
LN907888.1  241 TTACTTCCATTGTAGGTAACGTATTTGGTTTCAAAGCCCTACGTGCTCTACGTTTGGAGG
KR736525.1  240 TTACTTCCATTGTGGGTAACGTGTTTGGTTTCAAAGCCCTACGCGCTCTACGTTTGGAGG
KY627310.1  240 TTACTTCCATTGTGGGTAACGTATTTGGTTTCAAAGCCCTACGCGCTCTACGTTTGGAAG
KC483293.1  240 TTACTTCCATCGTGGGTAATGTATTTGGTTTCAAAGCGCTTCGCGCTCTACGTCTAGAGG
KJ204348.1  241 TTACTTCCATTGTGGGTAATGTATTTGGGTTCAAAGCCCTTCGCGCTCTGCGTCTGGAGG
KT458050.1  241 TTACTTCCATTGTGGGTAATGTATTTGGGTTCAAAGCCCTGCGTGCTCTACGTCTGGAGG
M77031.1    241 TTACTTCCATTGTGGGCAATGTATTTGGGTTCAAAGCCCTACGAGCTCTACGTCTGGAGG
AY875228.1  241 TTACTTCCATTGTGGGTAATGTATTTGGGTTCAAAGCCCTGCGTGCTCTACGTTTGGAGG
DQ907755.1  241 TTACTTCCATTGTGGGTAATGTATTTGGGTTCAAAGCCTTGCGTGCTCTACGTTTGGAGG
JQ933274.1  241 TTACTTCCATTGTGGGTAATGTATTTGGGTTCAAAGCCTTGCGTGCTCTACGTTTGGAGG
EF178544.1  241 TTACTTCTATTGTAGGTAATGTATTTGGTTTCAAAGCCCTACGAGCTCTACGCTTGGAAG
AB586281.1  241 TTACTTCTATTGTAGGTAATGTATTTGGTTTCAAAGCCTTACGAGCTCTACGCTTGGAAG
AM999843.1  241 TTACTTCTATTGTAGGTAATGTATTTGGTTTCAAAGCCCTACGAGCTCTACGCTTGGAAG
AB872559.1  241 TTACTTCCATTGTGGGTAATGTATTTGGGTTTAAAGCCCTGCGCGCTCTACGTCTAGAGG
EU643727.1  241 TTACTTCCATTGTGGGTAATGTATTTGGATTTAAAGCACTGCGTGCTCTACGTCTCGAAG
AY298820.1  241 TTACTTCCATTGTGGGTAATGTATTTGGTTTCAAAGCCCTACGAGCTCTACGTCTGGAGG
HM640507.1  241 TTACTTCCATTGTGGGTAATGTATTTGGTTTCAAAGCCCTACGAGCTCTACGTCTGGAGG
KC704911.1  241 TTACTTCCATTGTGGGTAATGTATTTGGTTTCAAAGCCCTGCGAGCTCTACGTCTGGAAG
KU748062.1  240 TTACTTCCATTGTGGGTAATGTATTTGGTTTCAAAGCCCTGCGAGCTCTACGTCTGGAAG
DQ875838.1  241 TTACTTCGATTGTGGGTAATGTATTTGGGTTCAAAGCTCTACGCGCTCTACGTCTGGAGG
EU840324.1  241 TTACTTCCATTGTGGGTAATGTATTTGGGTTCAAAGCCCTGCGTGCTCTACGTTTGGAGG
JX571893.1  241 TTACTTCCATTGTGGGTAATGTATTTGGCTTCAAAGCCCTGCGTGCTCTACGTTTGGAGG
AB586309.1  241 TTACTTCGATTGTGGGTAATGTATTTGGGTTCAAAGCCTTGCGTGCTCTACGTCTGGAAG
KX397733.1  240 TTACTTCCATTGTTGGTAATGTATTTGGGTTCAAGGCCCTGCGCGCTCTACGTCTGGAGG
JF944669.1  241 TTACTTCCATTGTGGGTAATGTATTTGGGTTCAAAGCTCTGCGCGCTCTACGTCTGGAGG
DQ535789.1  241 TTACTTCCATTGTGGGTAATGTATTTGGGTTCAAGGCCCTGCGCGCTCTACGTCTGGAGG
KM606572.1  241 TTACTTCCATTGTGGGTAATGTATTTGGGTTCAAAGCCCTGCGCGCTCTACGTCTGGAAG
HQ260780.1  239 TTACTTCCATTGTGGGTAATGTTTTTGGGTTCAAAGCGCTACGCGCTCTACGTCTGGAAG
MF065382.1  197 TTACTTCCATTGTAGGAAATGTATTTGGATTCAAAGCCCTACGTGCTCTACGTCTGGAAG
AY570449.1  241 TTACTTCCATTGTAGGAAATGTATTTGGATTCAAAGCCCTACGTGCTCTACGTCTGGAAG
AF190445.1  241 TTACTTCCATNGTAGGTAATGTATTTGGGTTCAAAGCCCTGCGGCCTCTGCGTCTGGAAG
HM849885.1  239 TTACTTCCATTGTAGGTAACGTATTTGGTTTCAAAGCCCTGCGTGCTCTACGTCTGGAAG
MG221719.1  240 TTACTTCCATCGTAGGTAATGTATTTGGGTTCAAAGCCCTGCGTGCTCTACGTCTGGAAG
JQ933508.1  241 TTACTTCCATTGTAGGTAATGTATTTGGGTTCAAAGCCCTGCGCGCTCTACGTCTGGAAG
JF308648.1  241 TTACTTCCATTGTAGGTAATGTATTTGGGTTCAAAGCCCTACGTGCTCTACGTCTGGAAG
AB744227.1  241 TTACTTCCATTGTAGGTAATGTATTTGGGTTCAAAGCCCTGCGCGCTCTACGTCTGGAAG
AM117226.1  241 TTACTTCCATTGTAGGTAATGTATTTGGGTTCAAAGCCCTGCGCGCTYTACGTCTGGAAG
```

```
AF419078.1   301 ATCTGCGAATTCCTCCCGCTTATTCCAAGACCTTTCAGGGTCCGCCCCATGGTATCCAGG
L34816.1     299 ATCTACGTATTTCTCCTGCTTACTGTAAGACTTTCTTAGGTCCGCCACACGGTATTCAGG
KP686040.1   301 ATTTACGGATTTCACCAGCTTACGTTAAAACTTTTCAGGGACCTCCTCATGGAATTGAAG
KJ446802.1   299 ATCTTCGTATTCCACCAGCATACGTAAAAACTTTCCAAGGGCCTCCTCATGGTATTCAAG
FN432103.1   301 ATTTAAGAATTCCAGCAGCTTATGCAAAAACTTTCCAAGGACCTCCTCATGGTATTCAGG
KX656068.1   299 ACCTTCGAATCCCTCCTGCTTATTCTAAAACTTTCATCGGACCGCCTCATGGTATTCAGG
JN572331.1   299 ACATTCGAGTTCCTCCTGCTTATTCTAAAACTTTTATTGGACCACCGCATGGCATTCAGG
U05608.1     301 ATATAAGGGTTCCACCTGCTTATTCTAAAACATTCTCAGGACCACCTCATGGTATCCAAG
KY089068.1   299 ATTTACGTATTCCTCCAGCTTATTCCAAAACTTTCCAAGGCCCACCTCATGGTATTCAAG
AY452692.1   299 ATTTACGTATTCCTCCAGCTTATTCCAAAACTTTCCAAGGTCCACCTCATGGTATTCAAG
JF940531.1   301 ATTTGCGGATTCCCCCTGCTTATTCCAAAACTTTTCAAGGTCCACCTCATGGTATCCAAG
GQ408938.1   299 ATCTTCGCATCCCCCCTGCTTATTCTAAAACTTTTCAAGGCCCGCCTCATGGTATCCAAG
HQ901564.1   301 ATTTGCGAATTCCTCCTGCTTATTCCAAAACTTTCCAAGGTCCACCTCATGGTATCCAGG
LN907888.1   301 ATCTACGAATTCCCACTGCTTATTCAAAAACTTTCCAAGGTCCGCCTCATGGTATCCAAG
KR736525.1   300 ATCTACGAATTCCCCCTACTTATTCAAAAACTTTCCAAGGTCCGCCTCACGGTATCCAAG
KY627310.1   300 ATCTACGAATTCCCACTAGTTATTCAAAAACTTTCCAAGGTCCGCCTCACGGTATCCAAG
KC483293.1   300 ATCTTCGAATTCCTCCTGCTTATGTTAAAACTTTCCAAGGCCCACCTCACGGTATCCAAG
KJ204348.1   301 ATCTGCGAATCCCTACTGCTTATGTGAAAACTTTCCAGGGCCCGCCTCACGGCATCCAAG
KT458050.1   301 ATCTGAGAATCCCTACTGCATATGTTAAAACTTTCCAAGGCCCGCCTCACGGCATCCAAG
M77031.1     301 ATCTGAGAATTCCTCCTGCTTATTCTAAAACTTTCCAGGGCCCACCTCATGGAATCCAAG
AY875228.1   301 ATTTGCGAATCCCTGTTGCTTATATAAAAACTTTCCAAGGCCCGCCTCACGGTATCCAAG
DQ907755.1   301 ATTTGCGAATCCCTGTTGCTTATATAAAAACTTTCCTAGGCCCGCCTCACGGTATCCAAG
JQ933274.1   301 ATTTGCGAATCCCTGTTGCTTATATAAAAACTTTCCTAGGCCCGCCTCACGGTATCCAAG
EF178544.1   301 ATTTGCGAATCCCCCCCGCTTATTCAAAAACTTTTCAAGGCCCACCTCATGGCATCCAAT
AB586281.1   301 ACTTACGAATTCCTCCTGCTTATGCAAAAACTTTCCAAGGTCCACCTCACGGTATCCAAG
AM999843.1   301 ACTTACGAATTCCCCCTGCTTATTCAAAAACTTTCCAAGGCCCGCCTCATGGTATCCAAT
AB872559.1   301 ATCTACGAATCCCTCCCGCGTATTCGAAAACTTTCCAAGGCCCCCCTCATGGCATCCAAG
EU643727.1   301 ATTTGCGAATCCCACCTGCGTATATTAAAACATTCCAAGGCCCGCCTCACGGCATTCAAG
AY298820.1   301 ATCTGCGAATTCCCCCTTCTTATTCAAAAACTTTCCAAGGCCCGCCTCATGGCATCCAAG
HM640507.1   301 ATCTGCGAATTCCCCCTTCTTATTCCAAAACTTTCCAAGGCCCGCCCCATGGCATCCAAG
KC704911.1   301 ATCTGCGAATTCCCACTTCTTATTCCAAAACTTTCCAAGGTCCGCCTCATGGCATCCAAG
KU748062.1   300 ATCTGCGAATTCCCACTTCTTATTCCAAAACTTTCCAAGGTCCGCCTCATGGCATCCAAG
DQ875838.1   301 ATCTGCGAATCCCTACCGCTTATGTTAAAACTTTCCAAGGCCCACCTCATGGTATCCAAG
EU840324.1   301 ATTTGCGAATCCCTCCTGCTTATACGAAAACTTTCCAAGGCCCGCCTCATGGTATCCAAG
JX571893.1   301 ATTTGCGAATCCCTCCTGCTTATTCGAAAACTTTCCAAGGCCCACCTCACGGTATCCAAG
AB586309.1   301 ATTTGCGAATCCCCCCTTCTTATTCTAAAACTTTCCAAGGCCCGCCTCACGGCATCCAAG
KX397733.1   300 ATTTGCGAATCCCTACTTCTTATACTAAAACTTTCCAAGGCCCGCCTCACGGCATCCAAG
JF944669.1   301 ATTTGCGAATCCCTACTGCTTATATTAAAACTTTCCAAGGCCCGCCTCATGGTATCCAAG
DQ535789.1   301 ATTTGCGAATCCCTCCTGCTTATACTAAAACTTTCCAAGGCCCGCCTCATGGTATCCAAG
KM606572.1   301 ATCTACGAATCCCTGCAGCGTATGCTAAAACGTTCCAAGGACCGCCTCATGGTATCCAAG
HQ260780.1   299 ATCTGCGAGTTCCTACTGCTTATATTAAAACTTTCCAAGGCCCGCCTCATGGCATCCAAG
MF065382.1   257 ATCTGCGAATCCCTGTTGCTTATGTTAAAACTTTCCAAGGCCCGCCTCATGGGATCCAAT
AY570449.1   301 ATCTGCGAATTCCTGTTGCTTATGTTAAAACTTTCCAAGGCCCGCCTCATGGGATCCAAG
AF190445.1   301 ATTTGCGAATTCCCATTTCTTATGTTAAAACCTTCCAAGGCCCGCCTCATGGCATTCAAG
HM849885.1   299 ATTTGCGAATTCCTACTGCGTATGTTAAAACTTTCCAAGGTCCACCTCACGGTATCCAAG
MG221719.1   300 ATTTGCGAATCCCTACTGCGTATGTTAAAACTTTCCAAGGTCCGCCTCACGGCATCCAAG
JQ933508.1   301 ATCTGCGAATCCCCGTTGCTTATGTTAAAACTTTCCAAGGACCGCCTCATGGCATCCAAG
JF308648.1   301 ATCTGCGAATTCCTACTGCTTATACTAAAACTTTCCAAGGCCCGCCTCATGGCATCCAAG
AB744227.1   301 ATCTGCGAATCCCTCCTGCTTATATTAAAACTTTCCAAGGTCCACCTCATGGGATCCAAG
AM117226.1   301 ATTTGCGAATCCCTCCTGCTTATATTAAAACCTTCCAAGGCCCGCCTCATGGCATCCAAG
```

```
AF419078.1   361 TCGAAAGGGATAAATTGAACAAGTACGGCCGTCCCTTGCTGGGATGTACCATCAAACCCA
L34816.1     359 TTGAACGTGATCGTCTGAACAAGTACGGTCGTCCGCTTCTAGGTTGTACAATTAAGCCTA
KP686040.1   361 TTGAGAGAGATAAATTAAATAAATATGGACGTCCTTTATTAGGATGTACTATTAAACCAA
KJ446802.1   359 TAGAACGTGATGAACTTAACAAATATGGTCGTGGTTTATTAGGTTGTACAATTAAACCAA
FN432103.1   361 TAGAAAGAGACAAACTAAATAAATATGGTCGTCCTCTTTTGGGTTGTACTATTAAACCAA
KX656068.1   359 TCGAGAGGGATAAACTGAACAAATATGGACGTCCTTTATTGGGATGTACAATCAAGCCAA
JN572331.1   359 TCGAAAGGGATAAACTGAACAAATATGGACGTCCCTTATTAGGATGTACAATCAAACCAA
U05608.1     361 TTGAAAGAGATAAACTGAACAAGTATGGTCGTCCTTTTTTAGGATGTACAATCAAACCAA
KY089068.1   359 TTGAAAGAGATAAATTAAATAAATATGGTCGTCCATTATTAGGATGTACTATTAAGCCAA
AY452692.1   359 TTGAACGAGATAAATTAAACAAATATGGTCGTCCATTATTAGGCTGTACTATTAAACCAA
JF940531.1   361 TTGAAAGAGATAAATTGAACAAATATGGCCGTCCTTTGTTGGGATGTACTATCAAACCAA
GQ408938.1   359 TAGAAAGAGATAAGCTGAATAAGTACGGCCGTCCTTTATTGGGATGTACTATTAAACCTA
HQ901564.1   361 TTGAAAGAGATAAATTGAACAAATACGGCCGTCCCCTATTGGGATGTACTATTAAACCAA
LN907888.1   361 TTGAAAGGGATAAGTTGAACAAGTATGGTCGTCCTTTATTAGGATGTACTATTAAACCAA
KR736525.1   360 TTGAAAGGGATAAGTTGAACAAGTATGGTCGTCCTTTATTGGGATGTACTATTAAACCAA
KY627310.1   360 TTGAAAGGGATAAGTTGAACAAGTATGGTCGTCCTTTATTGGGATGTACTATTAAACCAA
KC483293.1   360 TTGAAAGAGATAAATTGAATAAGTATGGTCGTCCCCTATTGGGATGTACTATTAAACCAA
KJ204348.1   361 TTGAGAGAGATAAATTGAACAAGTATGGACGTCCCCTATTGGGATGTACTATTAAACCGA
KT458050.1   361 TTGAGAGAGATAAGTTGAACAAGTATGGCCGTCCCCTATTGGGATGTACCATTAAACCGA
M77031.1     361 TTGAGAGAGATAAATTGAACAAGTATGGTCGTCCCCTATTGGGATGCACTATTAAACCCA
AY875228.1   361 TTGAGAGAGATAAATTGAACAAGTACGGCCGTCCTCTACTGGGATGCACTATTAAGCCGA
DQ907755.1   361 TTGAGAGAGATAAATTGAACAAGTACGGCCGTCCCCTATTGGGATGCACTATTAAACCTA
JQ933274.1   361 TTGAGAGAGATAAATTGAACAAGTATGGCCGTCCCCTATTGGGATGCACTATTAAACCGA
EF178544.1   361 CTGAAAGAGATAAGTTGAACAAGTACGGTCGTCCTCTATTGGGATGTACTATTAAACCAA
AB586281.1   361 CTGAAAGAGATAAGTTGAACAAGTATGGTCGTCCTCTATTGGGATGTACTATTAAACCAA
AM999843.1   361 CTGAAAGAGATAAGTTAAACAAATATGGTCGTCCTCTATTGGGATGTACTATTAAACCAA
AB872559.1   361 TTGAGAGAGATAAATTGAACAAGTATGGGCGCCCCCTATTGGGATGTACTATTAAACCTA
EU643727.1   361 TTGAGAGAGATAAATTGAACAAGTATGGTCGTCCCCTGTTGGGATGTACTATTAAACCAA
AY298820.1   361 TTGAAAGAGATAAGTTGAACAAGTATGGTCGTCCTCTATTGGGATGTACTATTAAACCAA
HM640507.1   361 TTGAAAGAGATAAATTGAACAAGTACGGTCGTCCCTTATTGGGATGTACTATTAAACCAA
KC704911.1   361 TTGAAAGAGATAAATTGAACAAGTATGGTCGTCCCCTATTGGGATGTACAATTAAACCAA
KU748062.1   360 TTGAAAGAGATAAATTGAACAAGTATGGTCGTCCCCTATTGGGATGTACTATTAAACCAA
DQ875838.1   361 TTGAAAGGGATAAATTGAACAAATATGGTCGACCCCTATTGGGATGTACTATTAAACCAA
EU840324.1   361 TTGAAAGAGATAAATTGAACAAATATGGACGTCCCCTATTGGGATGTACTATTAAACCGA
JX571893.1   361 TTGAAAGAGATAAATTGAACAAGTATGGGCGTCCCCTATTGGGATGCACTATTAAACCTA
AB586309.1   361 TTGAGAGAGATAAATTGAACAAGTACGGCCGTCCCCTATTGGGATGTACTATTAAACCTA
KX397733.1   360 TTGAGAGAGATAAATTGAACAAGTACGGCCGTCCCCTATTGGGATGTACTATTAAACCTA
JF944669.1   361 TTGAAAGAGATAAATTGAACAAGTATGGCCGCCCTCTATTGGGATGTACTATTAAACCAA
DQ535789.1   361 TTGAGAGAGATAAATTGAACAAGTATGGCCGTCCCCTATTGGGATGCACTATTAAACCAA
KM606572.1   361 TTGAAAGAGATAAATTGAACAAGTACGGTCGTCCCCTGTTGGGATGTACTATTAAACCTA
HQ260780.1   359 TTGAGAGAGATAAATTGAACAAGTATGGTCGTCCCCTACTGGGATGTACTATTAAACCAA
MF065382.1   317 CTGAGAGAGATAAATTAAACAAGTATGGTCGTCCCCTGTTGGGATGTACTATTAAACCTA
AY570449.1   361 TTGAGAGAGATAAATTGAACAAGTACGGTCGTCCTCTGCTGGGATGTACTATTAAACCTA
AF190445.1   361 TCGAGAGAGATAAATTGAACAAGTATGGTCGTCCCCTGTTGGGATGTACTATTAAACCTA
HM849885.1   359 TTGAAAGAGATAAATTGAACAAGTATGGTCGTCCTCTGTTGGGATGTACTATTAAACCTA
MG221719.1   360 TTGAGAGAGATAAATTGAACAAGTATGGTCGTCCCCTGTTGGGCTGTACTATTAAACCTA
JQ933508.1   361 TTGAGAGAGATAAATTGAACAAATATGGTCGTCCTCTGTTGGGATGTACTATTAAACCTA
JF308648.1   361 TTGAGAGAGATAAATTGAACAAGTATGGTCGTCCCCTGTTGGGATGTACTATTAAACCTA
AB744227.1   361 TTGAAAGAGATAAATTGAACAAGTATGGTCGTCCCCTGTTGGGATGTACTATTAAACCTA
AM117226.1   361 TTGAGAGAGATAAATTGAACAAGTATGGTCGTCCCCTGTTGGGATGTACTATTAAACCTA
```

```
AF419078.1   421 AACTAGGTCTGTCTGCCAAAAATTACGGCAGAGCCGTCTATGAATGTCTTCGTGGTGGAC
L34816.1     419 AACTAGGTCTTTCAGCTAAGAACTATGGTCGTGCTGTTTATGAATGTCTTCGTGGTGGTC
KP686040.1   421 AATTAGGTTTATCAGCTAAAAATTATGGACGAGCTGTTTATGAATGTTTACGAGGGGGAC
KJ446802.1   419 AATTAGGTCTTTCAGCGAAAAACTACGGTCGTGCTGTATACGAATGTTTACGTGGTGGTC
FN432103.1   421 AATTGGGTCTATCTGCAAAAAACTATGGTAGAGCTGTATATGAATGTTTACGTGGTGGAC
KX656068.1   419 AATTAGGTCTGTCTGCTAAGAATTATGGTAGAGCCGTCTATGAATGCCTTCGTGGTGGAC
JN572331.1   419 AATTAGGTCTGTCTGCTAAGAATTATGGTAGAGCCGTCTATGAATGCCTTCGTGGTGGAC
U05608.1     421 AATTGGGTTTATCGGCTAAGAATTATGGTAGAGCTGTTTATGAATGTCTCCGTGGGGGGC
KY089068.1   419 AATTAGGTTTATCTGCTAAAAACTATGGTAGAGCTGTATATGAATGTCTTCGTGGTGGAC
AY452692.1   419 AATTAGGTTTATCTGCTAAAAATTATGGTAGAGCTGTATATGAATGTCTTCGTGGTGGAC
JF940531.1   421 AATTGGGTCTATCGGCTAAGAACTATGGTAGAGCAGTTTACGAATGTCTTCGTGGTGGAC
GQ408938.1   419 AATTGGGATTATCCGCAAAGAACTATGGTAGAGCCGTTTATGAATGTCTGCGTGGTGGAC
HQ901564.1   421 AATTGGGATTATCCGCGAAAAACTACGGTAGAGCAGTTTATGAATGTCTTCGTGGTGGAC
LN907888.1   421 AATTGGGATTATCCGCAAAAAATTATGGTAGAGCGTGTTATGAGTGTCTACGCGGTGGAC
KR736525.1   420 AATTGGGATTATCCGC--------------------------------------------
KY627310.1   420 AATTGGGATTATCCGCAAAAAATTACGGTAGAGCGTGTTATGA-----------------
KC483293.1   420 AATTGGGTTTATCTGCTAAAAACTACGGTAGGGCGGTTTATGAAGTTCT-----------
KJ204348.1   421 AATTGGGATTGTCCGCTAAGAACTACGGTCGAGCAGTTTTTGAATGTCT-----------
KT458050.1   421 AATTAGGGTTATCCGCTAAGAACTACGGTAGAGCTGTTTATGAATGTCTTCGTGGTGGAC
M77031.1     421 AATTGGGGTTATCTGCAAAGAACTATGGTAGAGCCGTTTATGAGTGTCTCCGTGGTGGAC
AY875228.1   421 AATTGGGGTTATCTGCTAAAAACTATGGTCGAGCAGTTTATGAATGTCTTCGCGGTGGAC
DQ907755.1   421 AATTGGGGTTATCCGCTAAAAACTATGGTCGAGCAGTTTATGAATGTCTTCGCGGTGGAC
JQ933274.1   421 AATTGGGGTTATCCGCTAAAAACTATGGTCGAGCGGTTTATGAATGTCTTCGCGGTGGAC
EF178544.1   421 AATTAGGATTATCCGCAAAGAACTACGGTAGAGCATGTTATGAATGTCTACGCGGTGGAC
AB586281.1   421 AATTGGGATTATCCGCAAAGAATTACGGTAGAGCATGTTATGAATGTCTACGTGGTGGAC
AM999843.1   421 AATTGGGATTATCCGCAAAGAACTACGGTAGAGCATGTTATGAATGTCTACGTGGTGGAC
AB872559.1   421 AATTGGGATTATCCGCTAAGAACTATGGTAGAGCAGTTTATGAATGTCTACGTGGTGGAC
EU643727.1   421 AATTAGGGTTATCCGCTAAAAACTACGGCAGAGCAGTTTATGAATGTCTTCGTGGTGGAC
AY298820.1   421 AATTGGGATTATCCGCAAAGAACTACGGTAGAGCGGTTTATGAATGTCTACGCGGTGGGC
HM640507.1   421 AATTGGGATTATCCGCAAAAAACTACGGCAGAGCGGTTTATGAGTGTCTACGTGGTGGGC
KC704911.1   421 AATTGGGATTATCCGCAAAAAACTACGGTAGAGCGGTTTATGAATGTCTAAGGGGCGGAC
KU748062.1   420 AATTGGGATTATCCGCAAAAAACTACGGTAGAGCGGTTTATGAAT--------------
DQ875838.1   421 AATTGGGATTATCCGCTAAGAACTATGGTAGAGCGGTTTATGAATGTCTTCGCGGTGGAC
EU840324.1   421 AATTGGGGTTGTCCGCTAAGAACTACGGCCGAGCAGTTTATGAATGTCTTCGTGGCGGAC
JX571893.1   421 AATTGGGTTTATCTGCTAAGAACTACGGTCGAGCAGTTTATGAATGTCT-----------
AB586309.1   421 AATTGGGGTTATCCGCGAAGAATTACGGTAGAGCGGTTTATGAATGTCTCCGTGGTGGAC
KX397733.1   420 AATTGGGGTTATCTGCTAAGAATTACGGTAGGGCGGTTTATGAATGTCTC----------
JF944669.1   421 AATTGGGATTATCTGCTAAGAACTATGGTAGAGCAGTTTATGAATGTCTACGCGGTGGAC
DQ535789.1   421 AATTGGGATTATCCGCTAAGAATTATGGTAGAGCAGTTTATGAATGTCTACGCGGTGGAC
KM606572.1   421 AATTGGGGTTATCTGCTAAAAACTATGGGAGAGCTGTTTATGAATGTCTCCGCGGTGGAC
HQ260780.1   419 AATTGGGATTATCCGCTAAGAACTACGGTAGAGCGGTTTATGAATGTCTCCGCGGTGGAC
MF065382.1   377 AATTGGGGTTATCTGCTAAAAACTACGGTAGAGCATGTTATGAATGTCTTCGCGGTGGAC
AY570449.1   421 AATTGGGGTTATCTGCTAAAAACTATGGTAGAGCGGTTTATGAATGTCTTCGCGGTGGAC
AF190445.1   421 AATTAGGTTTATCTGCTAAAAACTATGGTAGAGCAGTTTATGAATGTCTTCGTGGTGGAC
HM849885.1   419 AATTGGGCTTATCCGCTAAAAACTACGGTAGAGCTGTTTATGAATGTCTTCGTGGTGGCC
MG221719.1   420 AATTGGGGTTATCCGCTAAAAACTACGGTAGAGCTGTTTATGAATGTCT-----------
JQ933508.1   421 AATTGGGGTTATCCGCTAAAAACTACGGTAGAGCGGTTTATGAATGTCTCCGCGGTGGAC
JF308648.1   421 AATTGGGGTTATCCGCTAAAAACTACGGTAGAGCTGTTTATGAATGTCTCCGCGGTGGGC
AB744227.1   421 AATTGGGGTTATCTGCTAAAAACTACGGTAGAGCTGTTTATGAATGTCTTCGCGGTGGAC
AM117226.1   421 AATTAGGTTTATCTGCTAAAAACTACGGTAGAGCTGTTTATGAATGTCTTCGCGGTGGAC
```

| | | |
|---|---|---|
| AF419078.1 | 481 | TCGATTTCAC |
| L34816.1 | 479 | TTGATTTCAC |
| KP686040.1 | 481 | TCGATTTTAC |
| KJ446802.1 | 479 | TTGATTTCAC |
| FN432103.1 | 481 | TTGATTTTAC |
| KX656068.1 | 479 | TTGATTTTAC |
| JN572331.1 | 479 | TTGATTTTAC |
| U05608.1 | 481 | TCGATTTTAC |
| KY089068.1 | 479 | TTGATTTCAC |
| AY452692.1 | 479 | TTGATTTCAC |
| JF940531.1 | 481 | TCGATTTTAC |
| GQ408938.1 | 479 | TTGATTTTAC |
| HQ901564.1 | 481 | TTGATTTTAC |
| LN907888.1 | 481 | TTGATTTTAC |
| KR736525.1 |  | ---------- |
| KY627310.1 |  | ---------- |
| KC483293.1 |  | ---------- |
| KJ204348.1 |  | ---------- |
| KT458050.1 | 481 | TTGATTTTAC |
| M77031.1 | 481 | TTGATTTTAC |
| AY875228.1 | 481 | TTGATTTTAC |
| DQ907755.1 | 481 | TTGATTTTAC |
| JQ933274.1 | 481 | TTGATTTTAC |
| EF178544.1 | 481 | TTGATTTTAC |
| AB586281.1 | 481 | TTGATTTTAC |
| AM999843.1 | 481 | TTGATTTTAC |
| AB872559.1 | 481 | TTGACTTTAC |
| EU643727.1 | 481 | TTGATTTTAC |
| AY298820.1 | 481 | TTGATTTTAC |
| HM640507.1 | 481 | TTGATTTTAC |
| KC704911.1 | 481 | TTGATTTTAC |
| KU748062.1 |  | ---------- |
| DQ875838.1 | 481 | TTGATTTTAC |
| EU840324.1 | 481 | TTGATTTTAC |
| JX571893.1 |  | ---------- |
| AB586309.1 | 481 | TTGATTTTAC |
| KX397733.1 |  | ---------- |
| JF944669.1 | 481 | TTGATTTTAC |
| DQ535789.1 | 481 | TTGATTTTAC |
| KM606572.1 | 481 | TTGATTTTAC |
| HQ260780.1 | 479 | TTGATTTTAC |
| MF065382.1 | 437 | TTGATTTTA- |
| AY570449.1 | 481 | TTGATTTTAC |
| AF190445.1 | 481 | TTGATTTTAC |
| HM849885.1 | 479 | TTGATTTTAC |
| MG221719.1 |  | ---------- |
| JQ933508.1 | 481 | TTGATTTTAC |
| JF308648.1 | 481 | TTGATTTTAC |
| AB744227.1 | 481 | TTGATTTTAC |
| AM117226.1 | 481 | TTGATTTTAC |

**Supplemental Figure S3:** Multiple sequence alignment of 50 sequences randomly drawn from the ITS2 database produced with Muscle and annotated with BoxShade (v3.21).

```
AJ293780.1    1 ATGCGATA---CGTAGTGTGAATTGCAGAAT-TCCGTGAACCATCGAATCTTTGAACGCA
HQ246426.1    1 ATGCGATA---CGTAGTGTGAATTGCAGAAT-TCCGTGAACCATCGAATCTTTGAACGCA
KC411873.1    1 ATGCGATA---CGTAATGTGAATTGCAGAAT-TCCGTGAATCATCGAATCTTTGAACGCA
MF034641.1    1 ATGCGATA---CGTAGTGTGAATTGCAGAAT-TCCGTGAATCATCGAATCTTTGAACGCA
JN631369.1    1 ATGCGATA---CCTAGTGTGAATTGCAGAAT-TCCGCGAATCATCGAGTCTTTGAACGCA
KP256679.1    1 ATGCGATA---CTTTGTGTGAATTGCAGAATCCCCGTGAATCATCGAATATTTGAACGCA
KX229790.1    1 ATGCGATA---TTTGGTGTGAATTGCAGAAT-CTCGTGAGCCATCAACTATTTGAATGCA
GQ866430.1    1 ATGCGATAGCGCGTGGTGTGAATTGCAGAAT-CCCGTGAACCATCGAGTTTTTGAACGCA
AJ489792.1    1 ATGCGATA---CCTGGTGTGAATTGCAGAAT-CCCGTGAACCATCGAGTTTTTGAACGCA
KT825439.1    1 ATGCGATA---CGTGGTGCGAATTGCAGAAT-CCCGCGAACCATCGAGAATTTGAACGCA
KP057199.1    1 ATGCGATA---CGTGGTGCGAATTGCAGAAT-CCCGCGAACCATCGAGAATTTGAACGCA
FJ213861.1    1 ATGCGATA---CGTGGTGTGAATTGCAGAAT-CCCGTGAACCATCGAATTTTTGAACGCA
KX087731.1    1 -------------------------------------------TCGAGTCTTTG-ACGCA
AF287681.1    1 ATGCGATA---CTTGGTGTGAATTGCAGAAT-CCCGTGAACCATCGAGTCTTTGAACGCA
GU572164.1    1 ATGCGATA---CTTGGTGTGAATTGCAGAAT-CCCGTGAACCATCGAGTCTTTGAACGCA
MG236967.1    1 -----------------------TGCAGAAT-CCCGTGAACCATCGAGTCTTTGAACGCA
AF202376.1    1 ATGCGATA---CTTGGTGTGAATTGCAGAAT-CTCGTGAACCATTGAGTCTTTGAACGCA
MG234931.1    1 -----------------------TGCAGAAT-CCCGTGAACCATCGAGTTTTTGAACGCA
FJ428091.1    1 ATGCGATA---CCTGGTGTGAATTGCAGAAT-CCCGTGAACCATCGAGTCTTTGAACGCA
KX873060.1    1 ATGCGATA---CCTGGTGTGAATTGCAGAAT-CCCGCGAACCATCGAGTCTTTGAACGCA
FN434493.1    1 ATGCGATA---CYTGGTGTGAATTGCAGAAT-CCCGCGAACCATCGAGTCTTTGAACGCA
JQ730037.1    1 ATGCGACA---CTTGGTGTGAATTGCAGAAT-CACGCGAACCATCAATTCTTTGAACGCA
KF196362.1    1 ATGCGATA---CTTGGTGTGAATTGCAGAAT-CCCGTGAACCATCGAGTCTTTGAACGCA
AM492068.1    1 ATGCGATA---CTTGGTGTGAATTGCAGAAT-CCCGTGAACCATCGAGTCTTTGAACGCA
JN999661.1    1 -----------------------TGCAGAAT-CCCGTGAACCATCGAGTCTTTGAACGCA
EU586943.1    1 ATGCGATA---CTTGGTGTGAATTGCAGAAT-CCCGTGAACCATCGAGTTTTTGAACGCA
KF677333.1    1 ATGCGATA---CTTGGTGTGAATTGCAGAAT-CCCGTGAATCATCGAGTTTTTGAACGCA
KP098091.1    1 ATGCGATA---CTTGGTGTGAATTGCAGAAT-CCCGTGAATCATCGAGTTTTTGAACGCA
KF498066.1    1 ATGCGACA---CTTGGTGTGAATTGCAGAAT-CCCGTGAACCATCGAGTCTTTGAACGCA
MF446079.1    1 ATGCGATA---CTTGGTGTGAATTGCAGAAT-CCCGTGAACCATCGAGTCTTTGAACGCA
HE586032.1    1 ATGCGATA---CTTGGTGTGAATTGCAGAAT-CCCGTGAACCATCGAGTCTTTGAACGCA
KJ845982.1    1 ATGCGATA---CTTGGTGTGAATTGCAGAAT-CCCGTGAACCATCGAGTCTTTGAACGCA
KM064970.1    1 -TGCGATA---CTTGGTGTGAATTGCAGAAT-CCCGTGAACCATCGAGTCTTTGAACGCA
JN253228.1    1 -----------------------TGCAGAAT-CCCGTGAACCATCGAGTCTTTGAACGCA
KY858482.1    1 ATGCGATA---CTTGGTGTGAATTGCAGAAT-CCCGTGAACCATCGAGTTTTTGAACGCA
AY625633.1    1 ATGCGATA---CTTGGTGTGAATTGCAGGAT-CCCGTGAACCATCGAGTCTTTGAACGCA
KX646449.1    1 ATGCGATA---CTTGGTGTGAATTGCAGAAT-CCCGTGAACCATCGAGTCTTTGAACGCA
MG236571.1    1 -----------------------TGCAGAAT-CCCGTGAACCATCGAGTCTTTGAACGCA
AY936715.1    1 ATGCGATA---CTTGGTGTGAATTGCAGAAT-CCCGTGAACCATCGAGTTTTTGAACGCA
MG219588.1    1 -----------------------TGCAGAAT-CCCGTGAACCATCGAGTTTTTGAACGCA
AY613151.1    1 --GCGATA---TTTGGTGTGAATTGCAGAAT-CCCGTGAACCATCGAGTTTTTGAACGCA
MG219618.1    1 -----------------------TGCAGAAT-CCCGTGAACCATCGAGTTTTTGAACGCA
MG519650.1    1 ATGCGATA---CTTGGTGTGAATTGCAGAAT-CCCGTGAACCATCGAGTTTTTGAACGCA
AB446176.1    1 --------------------AATTGCAGAAT-CCCGTGAACCATCGAGTCTTTGAACGCA
KU288713.1    1 ATGCGATA---CTTGGTGTGAATTGCAGAAT-CCCGTGAACCATCGAGTCTTTGAACGCA
KM037612.1    1 ATGCGATA---CTTGGTGTGAATTGCAGAAT-CCCGTGAACCATCGAGTCTTTGAACGCA
AY634813.1    1 ATGCGATA---CTTGGTGTGAATTGCAGAAT-CCCGTGAACCATCGAGTCTTTGAACGCA
EU528080.1    1 -TGCGATA---CTTGGTGTGAATTGCAGAAT-CCCGTGAACCATCGAGTCTTTGAACGCA
JQ255431.1    1 ATGCGATA---CTTGGTGTGAATTGCAGAAT-CCCATGAACCATCGAGTCTTTGAACGCA
AY233319.1    1 ATGCGATA---CTTGGTGTGAATTGCAGAAT-CCCGTGAACCATCGAGTCTTTGAACGCA
```

```
AJ293780.1    57 AATTGCGCTCGAGGCC---TCGGCCAAGAGCATGTCTGCCTCAGCGTCGG---------C
HQ246426.1    57 TATTGCGCCCAAGACC---TCGGTCGAGGGCATGTCTGCCTCAGCGTCG--GTATACACC
KC411873.1    57 TATTGCGGTCGAGGCT---TCGGCCGAGACCATGCCTGCCTCAGCGTCGTAAT---AGTG
MF034641.1    57 TATTGCGGTCGAGGCT---TCGGCCAAGACCATGTCTGCCTCAGCGTCGGTGT---GAAC
JN631369.1    57 AGTTGCGCCCGAGGCT---TGTCCCGAGGGCATGCCTGCCTGAGCGTCAC--------CG
KP256679.1    58 AGTTGCGCCCGAGGCC---TAGGTCGAGGGCACGTCTGGCTGGGTGTGAT-GAAAGATCG
KX229790.1    57 AGTTGAGCTCGAAGCC-GCTTGGCTGAGGGCACGCCTGCCTGGCGTCAT-TCCGAGCTA
GQ866430.1    60 AGTTGCGCCCGAGGCC-AATTGGCTAAGGGCACGTCCGCCTGGCGTCAA-GCGCGATTA
AJ489792.1    57 AGTTGCGCCTGAGGCC-TCTTGGTTGAGGGCACGTCTGCCTGGCGTCAC-GCCAAAAGA
KT825439.1    57 AGTTGCGCCCGAGGCC-AGCCGGCCAAGGGCACGTCCGCCTGGCGTCAA-GC---ATCG
KP057199.1    57 AGTTGCGCCCGAGGCC-AGCCGGCCRAGGGCACGTCCGCCTGGCGTCAA-GC---ATCG
FJ213861.1    57 AGTTGCGCCCGAGGCCTTTTTGGCCTAGGGCACGCCTGCCTGGCGTCA--GTTGCCCCT
KX087731.1    17 AGTTGCGCCCGAAACC-ATTAGGTTGAGGGCACGCCTGCCTGGGTGTCAC------ATGT
AF287681.1    57 AGTTGCGCCCGAAGCC-ATTAGGCTGAGGGCACGTCTGCCTGGGTGTCGC------CCAT
GU572164.1    57 AGTTGCGCCCGAAGCC-ATTAGGCTGAGGGCACGCCTGCCTGGGTGTCAC------ACGT
MG236967.1    37 AGTTGCGCCCGAAGCC-ATTAGGCCGAGGGCATGCCTGCCTGGGTGTCAC------ACAT
AF202376.1    57 AGTTGTGCCCGAGGCC-TTGTGGCCGAGGGCACGCCTGCTTGGCGTCATGGC-----AT
MG234931.1    37 AGTTGCGCCCGAAGCC-ATCTGGTTGAGGGCACGTCTGCCTGGCGTCAC------GTAC
FJ428091.1    57 AGTTGCGCCCGAGGCC-ATCCGGCTAAGGGCACGCCTGCCTGGCGTCAC-GC----TTT
KX873060.1    57 AGTTGCGCCCGAGGCC-ATTCGGCCGAGGGCACGCCTGCCTGGCGTCAC-GCAAAACA-
FN434493.1    57 AGTTGCGCCCGAGGCC-ACCCGGCCAAGGGCACGCCTGCCTGGCGTCAC-GCCAAAACA
JQ730037.1    57 AGTTGCGCCCGGAGCC-ATTCGGCCGAGGGCACGCCTGCCTGGGTGTCGT-GA---ATCC
KF196362.1    57 AGTTGCGCCCGAATCC-TTTCGGTCGAGGGCACGTCTGCCTGGGTGTCACGTA---CACA
AM492068.1    57 AGTTGCGCCCTAGGCC-ATTAGGCTGAGGGCACGCCTGCCTGGCGTCAC-GC---ATCG
JN999661.1    37 AGTTGCGCCCGAAGCC-TTTTGGCTGAGGGCACGTCTGCCTGGCGTCAC-GC---ATCG
EU586943.1    57 AGTTGCGCCCAAAGTC-TTTCGACCGAGGGCACGTCTGCCTGGGTGTCAC------GCAA
KF677333.1    57 AGTTGCGCCCAAAGTC-ATTAGGCTGAGGGCACGTCTGCCTGGGTGTCAC-GC---ATCG
KP098091.1    57 AGTTGCGCCCGAAGCC-ATCCGGCCAAGGGCACGTCTGCCTGGCGTCAC-GC---ATCT
KF498066.1    57 AGTTGCGCCCGATGTC-TTCAGGCTGAGGGCACGTCTGTTTGGCGTCAC-GC---ATGT
MF446079.1    57 AGTTGCGCCCGAAGCC-ATCAGGTTGAGGGCACGTCTGCCTGGCGTCAC-GC---ATCT
HE586032.1    57 AGTTGCGCCCGAAGCT---TCGGCCGAGGGCACGTCTGCCTGGCGTCAC-GC---ATCG
KJ845982.1    57 AGTTGCGCCCGAAGCC-ATCAGGTCGAGGGCACGTCTGCCTGGCGTCAC------ACGT
KM064970.1    56 AGTTGCGCCCGAAACC--TTTGGTCGAGGGCACGTTTGCCTGGGTGTCAC-AC---ATGG
JN253228.1    37 AGTTGCGCCCCAAGCC-ATTAGGCCGAGGGCACGTCTGCCTGGGTGTCAC------GCAT
KY858482.1    57 AGTTGCGCCCGAAGCC-ATTAGGTTGAGGGCACGTCCGCCTGGGTGTCAC-GC---ATTG
AY625633.1    57 AGTTGCGCCCGAAGCC-TTCTGGCCGAGGGCACGTCTGCCTGGGTGTCAC------GCAT
KX646449.1    57 AGTTGCGCCCCAAGCC-TTTTGGCCGAGGGCACGTCTGCCTGGGTGTCAC------AAAT
MG236571.1    37 AGTTGCGCCCCAAGCC-TTCTGGCCGAGGGCACGTCTGCCTGGCGTCAC------AAAT
AY936715.1    57 AGTTGCGCCCGAAGCC-TTTTGGTCGAGGGCACGTCTGCCTGGGTGTCAC------GCAA
MG219588.1    37 AGTTGCGCCCGAAGCC-ATTAGGCCGAGGGCACGTCTGCCTGGCGTCTC-AC---ATCG
AY613151.1    55 AGTTGCGCCCGAAGCC-ATTAGGCCGAGGGCACGTCTGCATGGCGTCAC-GC---ATCG
MG219618.1    37 AGTTGCGCCCGAAGCC-ATCCGGCCGAGGGCACGCCTGCCTGGCGTCAC-GC---ATCG
MG519650.1    57 AGTTGCGCCCGAAGCC-ATCCGGTTGAGGGCACGCCTGCCTGGCGTCAC-GC---ATCG
AB446176.1    40 AGTTGCGCCCGAAGCC-ATTAGGCCGAGGGCACGTCTGCCTGGCGTCAC-GC---ATCG
KU288713.1    57 AGTTGCGCCCGAAGCC-ATCAGGCCGAGGGCACGCCTGCCTGGCGTCAC------ACGC
KM037612.1    57 AGTTGCGCCCGAAGCC-ATTAGGCCGAGGGCACGCCTGCCTGGCGTCAC------ACGT
AY634813.1    57 AGTTGCGCCCGAAGCC-TTCAGGCCGAGGGCACGTCTGCCTGGCGTCAC------ATGT
EU528080.1    56 AGTTGCGCCCGAAGCC-ATTAGGCCGAGGGCACGTCTGCCTGGCGTCAC-GC---ATCG
JQ255431.1    57 AGTTGTGCCCGAAGCC-ATTAGGCCGAGGGCACGTCTGCCTGGCGTCAC-GC---ATAA
AY233319.1    57 AGTTGCGCCCGAAGCC-ATTAGGCCGAGGGCACGTCTGCCTGGCGTCAC-GC---ATCG
```

```
AJ293780.1  105 TACTA---CCCTCAACCAACTTTCCTATCGCAGGATAGGGAATG----------------
HQ246426.1  112 CCTCG---CCTCCCTCCCCTGTTGGAGTCGGCTCAGCGCTTGGCCGCTGGGCTT------
KC411873.1  111 CAACT---CCTCACCCCCTGCCCCTTGTGGTGGAC-------------------------
MF034641.1  111 CCTCA---CTCCCCCTCACGGG--------------------------------------
JN631369.1  106 CACCC---CCACCTACCCAATACTTTGTCGTGAGGGTTG---------------------
KP256679.1  114 TAGGG---CGCGCCCTCTTCGCCCCGTGCGAGGG--------------------------
KX229790.1  115 CGCTC---CTCCCCACCCCGTGTCCGGCTCGACTGCCGTTCTCGCGGATCAGGGGAGA--
GQ866430.1  118 CGTCGCTTCACGCTGCACCGGCCACCGACGAGGTGCGACGGTGTCGGCTC----------
AJ489792.1  115 CA------CTCCCTTGCAATCCTTGTGTG-------------------------------
KT825439.1  112 CGTCG---CTCCGTGCCAGCTCCGGCCCACCCAGCGGGTGGCGTCGGCCGAGGCCC----
KP057199.1  112 CGTCG---CTCCGTGCCAGCTCCGGCCCACCCAGCGGGTGGCGTCGGCCGAGGCCC----
FJ213861.1  115 CGGCG---CTCCATGCCATGTGTGATCTCTCACGGCT-----------------------
KX087731.1   70 CGTCA---CTCCCTATGTATTTGCTCAT--------------------------------
AF287681.1  110 CGTTG---CCCCAGTGCCCTGGCCTTGCGCTAGGCACCGAG-------------------
GU572164.1  110 CGTTA---CCCTCACGCAAATGTCTCTTGTTAGCCACAGTG-------------------
MG236967.1   90 CGTTA---CCCCCACGCACAGGTGCGCTGCTGGCCTGTGCG-------------------
AF202376.1  111 CGTCG---CTTTCGCTCCACGCATTGTTGGCG----------------------------
MG234931.1   90 ACACG---CCTCCCACAAAACCTTTCTCCATTTTTGGACAAGGATTTTGTG---------
FJ428091.1  111 CGACG---CTTCGTCGTTGCACCCTCGGGGGGTGGGGGC---------------------
KX873060.1  114 ---CG---CTCCCACCCCTCATCGAGGAGCG----------------------------
FN434493.1  115 CG------CTCCCACTCCCCCTCACGGGGCGGAG--------------------------
JQ730037.1  112 ACTTG---TCCCCCGGCTCCCGCGGGCGCCCATGGGCGCGGGGATATGCCG---------
KF196362.1  113 CGTCT---CTCACAAACCTTTCCCTTCATGGGCGAAGGGTCTTGC---------------
AM492068.1  112 TGTCG---CTCCCCCCAGCCACCCCATGGGATGGCCGGTTT-------------------
JN999661.1   92 CGTCT---CCCCACATCCATCCCACACTAGTGCGGGGGA---------------------
EU586943.1  110 CATTG---CTCCCAACCTATTA--------------------------------------
KF677333.1  112 CGTCG---CCTCCTGAATAAAAATATA---------------------------------
KP098091.1  112 CGTCG---CCACCCCCTCCAGACCCGTCCTCCGGGACGCTGGCTGGCAGCGG--------
KF498066.1  112 CGTCAACCCTTCCCACGTCGTCTTCTTCCCCACTCAAAGGTGTCGGGGGACGATCTTACG
MF446079.1  112 CGTCG---CCCCCTCCACACGTCACTCAGAGTGCATGCGAAGG-----------------
HE586032.1  110 CGT-----CTCCCACACACACCCTGTGTGTCGT---------------------------
KJ845982.1  110 CGTTGCTCCCCCCCAAACCCCCCTCCCGCTCCTAGCCGGGCGAGGGGGGACCGTG-----
KM064970.1  110 CGTTG---CCCCCAATCCCCTCCGCCCTCTCAACGGGGCGAGCGGGACTC----------
JN253228.1   90 CGTCG---CCCCCATCCAACCCTGAGCCCTCGGGCCTCGGCTGGACCG------------
KY858482.1  112 CGTCG---CCCCCTCTCATCCCTCTTTACCAATGTTAAAGTCGGGGTGGAGTTGA-----
AY625633.1  110 CGTCG---CCCCCCAAACCCCCCAAACACAAGGGTGGGGAGGAATTGT------------
KX646449.1  110 CGTCG---TCCCCCATCCTCTTAAGGATA-------------------------------
MG236571.1   90 CGTCG---TTCCCCTCACAAAATTATGCGAGTG---------------------------
AY936715.1  110 CGTCG---CTCCCTCACTCCCTCACTCGAGGCATGTGAATT-------------------
MG219588.1   92 CGTCG---CCCCCAACCTCATTCCCTTCAGGAATGACGGTCTT-----------------
AY613151.1  110 CGTCA---CCCCCCTCCTACTCCCTCGGGACGCTGGGGT---------------------
MG219618.1   92 CGTCG---CCCCCACCAAACTTCCCCTTGGGGATACATGGCAT-----------------
MG519650.1  112 CGTCG---CCCCCGTCATAGTTCCCTTAAGGGTAGTCGTGGTGATT--------------
AB446176.1   95 CGTCG---CTCCCCCACCCGCGTTGGCGAG------------------------------
KU288713.1  110 CGTTG---CCCCCCCGCGCATCCCTCGGGAGCGTCG------------------------
KM037612.1  110 CGTTG---CCCCCCCCCAAACCCCTCGGGAGTTGGG------------------------
AY634813.1  110 CGTTGGACCCCCGACCCCTCCGTGGGTCGGA-----------------------------
EU528080.1  111 CGTCG---CCCCCAACCCATGTCCTCCGTGTCGCACGGGTGACCGGGA------------
JQ255431.1  112 CGTCA---CCCCCCGCACGCCGCAGGGCGTCG----------------------------
AY233319.1  112 CGTCG---CCCCCCAACCCCGCACTCCCTCATGGGGGTCGAGGCAG--------------
```

```
AJ293780.1  146 ---CCGGTTGGATA--TGGCCGTCC--------GG--------------TCTGCCCTTCG
HQ246426.1  163 ---GGAGGCGGATG--TGGCTTTCCCACGACTTGT-----------------GCG-----
KC411873.1  143 ---GGGAGTGGACC--TGGTCTCCT--------CA-----------------TCGCCTGG
MF034641.1  130 ---TGGAGTGGACC--TGGCCTCCTCAGTCCTAGT-----------------GCC-----
JN631369.1  142 ---GGATGTGTGGAAGTGGCTGTCC--------AT-----------------GACCCCGG
KP256679.1  145 ---GGCAGCGGATG--TGGTCGTCC--------GT-----------------GTCTCCCG
KX229790.1  170 ---GGAAGCGGACGT-TGGCCCCCC---GATCCGC-----------------GCGTGCGG
GQ866430.1  168 ---GAACGCGGAGAG-TGGCCCTTC--------GC-----------------GCAGATCT
AJ489792.1  138 ---GGATGTGGTGTT-TGGCCTTCC--------AT-----------------TCTACTTR
KT825439.1  165 ---GGATGTGCAGAG-TGGCCCGTC--------GT-----------------GCCCGTCG
KP057199.1  165 ---GGATGTGCAGAG-TGGCCCGTC--------GT-----------------GCCCGTCG
FJ213861.1  149 ---GGATGCGGGCC--TGGCCCTTC--------GA-----------------GTCCCTGA
KX087731.1   95 ---TGGGGTGAAGGT-TGGCTTCCT--------GC-----------------GAGACTAA
AF287681.1  148 ---CGGGGCGAGTGC-TGGCTTCCC--------GT-----------------GAGCAACG
GU572164.1  148 ---TAGGGTGTATGC-TGACCTCCC--------GC-----------------GAGCGGTG
MG236967.1  128 ---TGTAGTGTATGC-TGGCCTCCC--------GC-----------------GAGCGGTG
AF202376.1  140 ---GCGAGCGGAAAT-TGGCCCCGT--------GT-----------------GTCCTCGG
MG234931.1  138 ---GCGAGCAGAGAG-TGGCCTCCC--------GT-----------------GCAACTTT
FJ428091.1  147 ---GAACGCGGAGGA-TGGCCCCCC--------GT-----------------GCCGAAAG
KX873060.1  140 ---GGACGCGGCATG-TGGCCCCCC--------GTC----------------GCGCAAGG
FN434493.1  143 ---GGACGCGGCGTC-TGGCTCCCC--------GT-----------------GCCGTCAG
JQ730037.1  160 ---GGGCGCAGAATT-TGGCCTCCC--------GTCGGATCGGTCGTGTGCGGCACGCCG
KF196362.1  155 ---GTGAGCGGAGTT-TGGTATCCC--------GT-----------------GCTTGTAT
AM492068.1  150 ---GGGAGCGGAGAT-TGGCCCCCC--------GT-----------------GTGTCGTA
JN999661.1  128 ---AGGGGAGGATGA-TGGCCTCCC--------AT-----------------GCCTCACC
EU586943.1  129 ---GGGTGCGAAATA-TGGCCTCCC--------GT-----------------GCGATTTC
KF677333.1  136 ---GGGAGCGGATTA-TGGCTTCCC--------GT-----------------GCTTTGAT
KP098091.1  161 ---GGCGGCGGATGC-TGGCCTCCC--------GT-----------------TCCCTCGC
KF498066.1  172 GAGGAGGGTGGGATATTGGCCTCCC--------GT-----------------TATCCTTG
MF446079.1  152 ---AGGGGCGGATAT-TGGCCTCCC--------GT-----------------TATCCTTG
HE586032.1  138 ---GTAGGAGGAAGA-TGGCTTCCC--------GT-----------------GCCTCACC
KJ845982.1  165 ---GGGGGCGGACGA-TGACCTCCC--------GT-----------------GCGCGCTT
KM064970.1  157 ---GGGCGCGTAGGA-TGGCCTCCC--------GC-----------------GACGACCA
JN253228.1  135 ---CGGGGCGGAAAT-TGGCCTCCC--------GT-----------------GCGCTCAC
KY858482.1  164 ---GGGGGCGGACAT-TGGCCTCCC--------GT-----------------GCTGTTCT
AY625633.1  155 ---TGGGGCGGATAC-TGGTCTCCC--------GT-----------------GCGCTCCC
KX646449.1  136 ---CGGGACGGAAGC-TGGTCTCCC--------GT-----------------GTTTTACC
MG236571.1  120 ---CGGGACGGAAGC-TGGTCTCCC--------GT-----------------GTGTTACC
AY936715.1  148 ---CGGGGCGGAAAA-TGGCCTCCC--------GT-----------------GAACTTCG
MG219588.1  132 ---GGGAGCGGAAAT-TGGTCTCCC--------GT-----------------GCACGNTG
AY613151.1  146 ---GGGGGCGTAAAT-TGGCCTCCC--------GT-----------------GCGTGCAA
MG219618.1  132 ---CGGGGCGGAGAT-TGGCCTCCC--------AT-----------------TCTTTTGG
MG519650.1  155 ---GGGAGCGGAAAT-TGGCCTCCC--------GT-----------------GCTTGTTG
AB446176.1  122 ---GGGGGCGGATAT-TGGCCTCCC--------GT-----------------GCGCCTCG
KU288713.1  143 ---GGGGGGGGACGA-TGGCCTCCC--------GT-----------------GCGTCACC
KM037612.1  143 ---CGGGACGGATGA-TGGCCTCCC--------GT-----------------GTGCTCTG
AY634813.1  141 ---CGGGACGGATGA-TGACCTCCC--------GT-----------------GTGCCCCG
EU528080.1  156 ---TGGGGCGGAAAC-TGGCTTCCC--------GT-----------------GCTTTCGT
JQ255431.1  141 ---TGGGGCGGATAC-TAGCCTCCC--------GT-----------------GCGCTTCG
AY233319.1  155 ---AGGGGCGGATAC-TGGCCTCCC--------GT-----------------GTCTCACC
```

```
AJ293780.1  179 GGTTG--------------CCGGTCAGCTGAAATACATTCACGCTGTGGGAGCTGCTTTC
HQ246426.1  196 -------------------TGGGTCGGCTGAAAAGCAGAGGCTAGAGCTAGGACCCATCA
KC411873.1  173 CGA----------------TGGGCTGGCTGAAACTCGGAGAT---------------TCA
MF034641.1  163 -------------------TGGGTTGGCTGAAATTCGGAGGC---------------TCC
JN631369.1  174 TGCCTGGATGCCGGTGGTGGTGGTCGGCTGAAATGGAGCATT-----------CTGGACG
KP256679.1  175 AGGT---------------GCGGTCGGCTGAAA---AGTTGCCGTCGATCGTGTTTTCTG
KX229790.1  206 T------------------TCGGTGGGCACAAG---TGCGGT-------------TTGCC
GQ866430.1  199 GCGCG--------------ACGCGGGGTTGAAG---ACCGGA---------------TTG
AJ489792.1  169 TAAGGT-------------GTGGTAGGCTAAAGGAAGTGG-----------------CTG
KT825439.1  196 GT-----------------GCGGCGGGTTGAAG---AGCGGG---------------TCA
KP057199.1  196 GT-----------------GCGGCGGGTTGAAG---AGCGGG---------------TCA
FJ213861.1  179 GC-----------------TCGGCGGGCCAAAGCGTTTGGCT---------------TGC
KX087731.1  126 TTCCT------------C-GTGGTTGGTTGAAA---ACTAGT---------------TTA
AF287681.1  179 CCTC---------------GCGGTTGGTTGAAA---TATGAG---------------TCC
GU572164.1  179 TCTC---------------GTGGTTGGTTTAAA---ACTGAG---------------TTT
MG236967.1  159 CCTC---------------GTGGTTGGTTGAAA---TTCGAG---------------TCC
AF202376.1  171 GC-----------------ACAGTCGGTCGAAG---AGTGGG------------------
MG234931.1  169 TGT----------------GCGGTTGACCCAAA---AAAGAG---------------TAC
FJ428091.1  178 GT-----------------GCGGTTGGCCGAAG---AGCGGG---------------CCG
KX873060.1  172 G------------------GCGGTGGGCCGAAG---ATATGG---------------CTG
FN434493.1  174 GC-----------------GCGGAGGGCCGAAG---ATCGGG---------------CTG
JQ730037.1  208 TGGT---------------GCGGTTGGCCGAAAGTCTAGCAT-----------------T
KF196362.1  186 TGT----------------GCGGCTTACCTAAAAATTGAGAG---------------CCA
AM492068.1  181 TGCGC------------ACGCAGCCGGCCTAAAA--GCCGGG---------------CCC
JN999661.1  159 GGGT---------------GTGGCTGGCCTAAAA--AAGGAG-----------------C
EU586943.1  160 CTCTC--------------GCGGTTGGCCAAAAAGAATGG-----------------TCC
KF677333.1  167 -------------------GCGGTTGGCCTAAA-----AGAT---------------TGC
KP098091.1  192 GGC----------------GCGGCTGGCCCAAATGCTTCGGC---------------CCC
KF498066.1  207 AGCGT------------A-GCGGCTGGCCCAAA---CAATAT---------------ACC
MF446079.1  183 TGTA---------------GCGGCCGGCCCAAA---TAACAT---------------ACC
HE586032.1  169 CGGC---------------GCGGTTGGCCTAAAT--TGGGAG---------------ACC
KJ845982.1  196 CGACCC-------------GCGGTTGGTCCAAAA--ATGGAG---------------TCC
KM064970.1  188 CGTC---------------CCGGTTGGCCCAAA---ATCGAG----------------CG
JN253228.1  166 AGCCA--------------GCGGTTGGCCTAAA---TTCGAG---------------TCC
KY858482.1  195 TGT----------------GTGGCTGGCCTAAA---TTTGAT---------------TCA
AY625633.1  186 GCTC---------------GCGGTTGGCCCAAA---TACCGG---------------TCC
KX646449.1  167 GAAT---------------GCGGTTGGCCAAAA---TCTGAG---------------CTA
MG236571.1  151 GCAC---------------GCGGTTGGCCAAAA---TCTGAG---------------CTG
AY936715.1  179 TGATT--------------GCGGTTGGCCCAAA---CATGAG---------------ACC
MG219588.1  163 CTGC---------------GCGGTTGGCCTAAA---TAAGAG---------------TCG
AY613151.1  177 GCGT---------------GCGGCCGGCCCAAA---TGCGAT---------------CCC
MG219618.1  163 T------------------GTGGTTGGCCTAAA---CCGGAG---------------TCC
MG519650.1  186 T------------------GCGGTTGGTCAAAA---TAGGAG---------------TCC
AB446176.1  153 GCGT---------------GCGGCCGGCCCAAA---TGCGAT---------------CCC
KU288713.1  174 CCGC---------------GCGGTTGGCACAAAT--GCCGGG---------------TCC
KM037612.1  174 TCAT---------------GCGGTTGGCATAAA---AACAAG---------------TCC
AY634813.1  172 TCAC---------------GCGGCTGGCATAAAT--ACCAAG---------------TCC
EU528080.1  187 -------------------GCGGCTGGCCTAAA---CACGAG---------------TCC
JQ255431.1  172 AGGCC--------------GCGGCCGGCCTAAA---TGCGAG---------------TCC
AY233319.1  186 AC-----------------GCGGTTGGCCCAAA---TGTGAG---------------TCC
```

```
AJ293780.1  225 CTTTCGAGGGAAGGCGAGTTTCGAT-----------------TTGGT-------AGGTGA
HQ246426.1  237 TGGGCTCAACTGGATAGGTAGCCAC-----G-----------GTGCT-------TCACGG
KC411873.1  202 CGCTCGGGCC-TGTACGGCAACAGCAAGGTAG----------GTGGC-------TCGTTC
MF034641.1  189 CGCTCGGGCC-TGTATGGCAGCAGCAAGGTAG----------GTAGT-------TTACTA
JN631369.1  223 TGGTCGGAGA-AGAGAGGCTGCGAGAAGGTGATCCCCCCCTTGTTGGGAGGAGTTGGACA
KP256679.1  217 GCTTCTACCC-GGAC--GGAGCGAT-GGATG-----------GTGGTGGTGGTCTGTGCG
KX229790.1  232 GCGGAGCCGT-GCATGGGTCGCGGCGATTCG-----------GTGGTACACACGTGACCG
GQ866430.1  227 CCTTCCTCTT-GGCCATGCTTTGATAAAGGG-----------GTGGA-------TGGGTG
AJ489792.1  199 CTGGCTATG--GTGCCAGTCACAGC-ACTTG-----------GTGGA-------TGACAC
KT825439.1  221 TCGTCTCAT--CGGCGACGAGCAGC-GAGGG-----------GTGGA-------TGAAAG
KP057199.1  221 TCGTCTCAT--CGGCGACGAGCAGC-GAGGG-----------GTGGA-------TGAAAG
FJ213861.1  207 CCATTG-----GGGCTGGCTGTGGC-GAGTG-----------GTGGA-------TCGATC
KX087731.1  155 GCGGTG-----CAGTCCCCTGCAAT-AAATG-----------GTGGA-------TGAGAA
AF287681.1  206 GTGGTGGA---GGGC--GCCGTGAT-GGATG-----------GTGGT-------TGAGTA
GU572164.1  206 GCGGTG-----GAGTGTGCCGTGATGAAATG-----------GTGGA-------TGGGCA
MG236967.1  186 GCGGTG-----GGGTGCGCCGCGAT-AAATG-----------GTGGA-------CGGGCA
AF202376.1  193 TAGTCGGCAG-TCGTCGGGCACGAT-GGGTG-----------TTGGTCGCCGCGAGCGGG
MG234931.1  195 CGAGTGAC----GAAGTGTCACGAT-AAGTG-----------GTGGA------CTGTAAG
FJ428091.1  203 TCGGTGG----TTGTCGAACACGAC-GCGTG-----------GTGGATGCCTTGTGCGAG
KX873060.1  196 CCGGCGTATC-GTGCCGGACACAGC-GCGTG-----------GTGGG-------CGACCT
FN434493.1  199 CCGGCGTACC-GTGTCGGGCACAGC-GCGTG-----------GTGGG-------------
JQ730037.1  236 GCGGCGAT---GGGC--TCCACGGC-GTTCG-----------GTGGATCGTATGCGATTG
KF196362.1  215 GTGATG-----AAAC--TTCACGTC-TAGTG-----------GTGGT-------TAAGGA
AM492068.1  212 CCGGCGGT---CGAC--GTCACGAC-GAGTG-----------GTGGTTGATACGTGATAC
JN999661.1  185 CTTGAGTCAT-GGACT-GCAGCGGC-GTATG-----------GTGGT------ATACAAG
EU586943.1  189 TCGGCAGC---GAAT--GCCTCGAC-AATCG-----------GTGGT-------TGTAAG
KF677333.1  188 TCGCTCGCCGCGCAC--ATTGCGAC-AAGTG-----------GTGGT-------CGATTA
KP098091.1  221 CGGACAAG---GGAC--GTCGCGAC-TCAAG-----------GTGGT-------TGGACA
KF498066.1  236 GTGCCGAAGG-AAGT--CGCACGAT-ACGTG-----------GTGGT-------TGGATT
MF446079.1  210 GTGTCGAC---GGATGTCACACGAT-GCGTG-----------GTGGT-------TGAATT
HE586032.1  197 ACGGCATT---GAGA--GTCGCGGC-ATTAG-----------GTGGA-------TTTCTA
KJ845982.1  226 CCTGTCACGTC-------GTCTTGGC-AACAG-----------GTAGT-------CGATCA
KM064970.1  214 TCGGAGCGAT-CAGC--ACCACGRC-ATTCG-----------GTGGT-------TGAYTA
JN253228.1  194 TCGACGTCAT-CATC--GTCGCGAC-CATCG-----------GTGGTAATGCTGCAAGCA
KY858482.1  221 CTGTCGAC---AAAT--GTCACAAC-TAGTG-----------GTGGT-------TGAATC
AY625633.1  213 CTGGCAACGG-TG----GCCATGGC-AAGCG-----------GTGGT-------TGAGAG
KX646449.1  194 AGGACGCC---AGGAGTGTCTCGAC-ATGCG-----------GTGGTGAA----TTCAAG
MG236571.1  178 AGGATGCTGG-GAGC--GTCCCGAC-ATGCG-----------GTGGTGA-----TCTAAA
AY936715.1  207 AATTCGGC---CAAT--GCCGTGGC-ATTCG-----------GTGGT-------TGAAAA
MG219588.1  190 ACGTGGAC---GAGC--GTCACAAC-AAGTG-----------GTGGT-------TGTCAA
AY613151.1  204 GCATCGAC---GGAT--GTCACGAC-CAGTG-----------GTGGT-------TGAAAC
MG219618.1  187 CCTTCGGT---GGAC---GCACGAC-TAGTG-----------GTGGT-------TGAAAA
MG519650.1  210 CCTTCGGT---GGAC---ACACGGC-TAGTG-----------GTGGT-------TGTAAA
AB446176.1  180 CCGGCGAC---TCGT--GTCGCGAC-AAGTG-----------GTGGT-------TGAACA
KU288713.1  202 TCCGCGAC---GAAC--GACACCAC-AATCG-----------GTGGT-------CGGCCA
KM037612.1  201 TCGGCGAC---TAAC--GCCACGAC-AATCG-----------GTGGT-------TGTCAA
AY634813.1  200 CCGGCGGC---CGAC--GCCACGAC-AATCG-----------GTGGT-------TGTGAG
EU528080.1  210 CTAGCGAC---GGAC--GTTGTGAC-AAGTG-----------GTGGT-------TGATTT
JQ255431.1  200 ACGTCGAC---GGAC--GTCGCGAC-AAGTG-----------GTGGT-------TGAAAG
AY233319.1  211 TTGGCGAC---GGAC--GTCACGAC-AAGTG-----------GTGGT-------TGTAAA
```

```
AJ293780.1  261 CCTT------------------------------------------------------TG
HQ246426.1  274 CACC----------------------------------GCCTACACGAAGTTGATGCTTG
KC411873.1  244 ACTC-----------------------------------------------CGGCTGATG
MF034641.1  231 CTCC------------------------------------------------GGCTGATG
JN631369.1  282 GCGC------------------------------CTCTCCCTCCGGGGATCTGTTGGATG
KP256679.1  262 TTCCCTGCGCTGAGCGCGCACGAGGTACGTTGGCCCCCTCCCCGCCCTTCTCCTTCGGTG
KX229790.1  280 CCGT-------------------------------------CCTGGCGCCTCCCCTGCCG
GQ866430.1  268 CTGC---------------------------------------CAGAGGGCCCGCTATCA
AJ489792.1  238 ATGT---------------------------------------------TGTTCTCGAAG
KT825439.1  260 TTGT-------------------------------------------------GCCTGTG
KP057199.1  260 TTGT-------------------------------------------------GCCTGTG
FJ213861.1  243 GCTC---------------------------------------AGACGAGCCGTACGTCG
KX087731.1  191 ACTC-----------------------------------------------CTCTCGAGA
AF287681.1  242 AAAG-------------------------------------------------CTCGAGA
GU572164.1  243 TTGC----------------------------------------------TCGAGGCCAG
MG236967.1  222 ATGC--------------------------------------------------CCGAGA
AF202376.1  240 AACA---------------------------------------------------GAACG
MG234931.1  233 CCCT-----------------------------------TAGTGGCTTTGCCAAATCGAG
FJ428091.1  247 CCGT-----------------------------------------------------ACG
KX873060.1  236 CGCT-----------------------------------------------TTACTTCCG
FN434493.1  233 -----------------------------------------------ATTCTACTCGGCG
JQ730037.1  279 GCTG------------------------------------------------------TG
KF196362.1  249 ACCT----------------------------------TCATGGGTCAAACCAGTTCGCG
AM492068.1  255 CCCT------------------------------------------TGCCTTATCGGACG
JN999661.1  225 GCTT---------------------------------TGCCTCGAATGCATCGTGCCGTG
EU586943.1  225 ACCT----------------------------------------------TCGGACACAG
KF677333.1  227 TATT---------------------------------------------GAACGCAAATG
KP098091.1  257 TCAT-----------------------------------------------------TCA
KF498066.1  274 CCTC-------------------------------------------AACTCGCAAGCTA
MF446079.1  248 CGTC-------------------------------------AACTTGCGAACTATCTACA
HE586032.1  233 TTCC-----------------------------------------------------ATG
KJ845982.1  261 TTCG-----------------------------------------------GTGCCACCG
KM064970.1  252 GACC-------------------------------------------CCAATGATCAATG
JN253228.1  239 AACC----------------------------------------------TCGTTCGGAG
KY858482.1  257 AATC------------------------------------------------GCGTGCTG
AY625633.1  249 ACCC---------------------------------------------TCGGATAAATG
KX646449.1  235 CCTC-----------------------------------------------TTTAGTTTG
MG236571.1  218 GCCT----------------------------------------------CTTCATATTG
AY936715.1  243 TACC-----------------------------------------------TTACTATTG
MG219588.1  226 ACCG---------------------------------------------TTGCGTTGCTG
AY613151.1  240 CTCA--------------------------------------------ACTTGCGTGCTG
MG219618.1  222 GACC---------------------------------------------CTCGTCCTGTG
MG519650.1  245 GACC---------------------------------------------CTTTTCTTCTG
AB446176.1  216 TCTC------------------------------------------AATCTCGCGTCTCG
KU288713.1  238 ACCT----------------------------------------------CGGTTGCCAG
KM037612.1  237 ACCT----------------------------------------------CTGTTGCCTA
AY634813.1  236 ACCT----------------------------------------------CGGTGTCCTG
EU528080.1  246 TCTC-------------------------------------------AACTAGAGCGCTG
JQ255431.1  236 CTCA-------------------------------------------ACTCTCTCTGCTG
AY233319.1  247 AAGC--------------------------------------------CCTCTTCTCATG
```

```
AJ293780.1  267 CCGGTTA--CATGCCTT--------------CGTTGCCCGCTAAGCTTCCCC--------
HQ246426.1  300 CGGTACTGGCTGGCGGCCTGCAGGAAACGTGTGGTATGCTTGGTGCAAAGCT--------
KC411873.1  257 CTTAGGG--CACGCG----------------CACGGATCCTTCAGAAAAACT--------
MF034641.1  243 CTTTGGG--CACGTAGC----------------ATGAGCCCG------------------
JN631369.1  312 TCGGGCT--CCTCTGGGGGCCCGGGAAAT--GGAGGGAGTCGGAGTGGCTCC--------
KP256679.1  322 CGATGCG--GGACCGAGCGAC----------CGGCGGGGCG--ATGTTCCTCCTTTGCG
KX229790.1  303 TCGCGCG--GCAGCTCT--------------TTGAGCCCTCTCCGGACGCCA--------
GQ866430.1  289 CTTCGTG--TCCCTGGCGAGGAAGACAGATACACACATCCCTGGTGGCTCAC--------
AJ489792.1  253 CTGTGACTTGTACTAATTCCGGTAA------CATTGGCCCTT--TCGACCCA--------
KT825439.1  271 CTGCGTTCGTGCCTG-----------------------CCCGAAAAAAGATT--------
KP057199.1  271 CTGCGTTCGTGCCTG-----------------------CCCGAGAAAAGATT--------
FJ213861.1  264 CCCAGCT--AGACCCCTCGGGT---------ATATGGCCTTGTTGAAACCCC--------
KX087731.1  204 CCAATTG--TGGGTGTGG-------------TACTGTGCTTGATGGAATCATTTGACCCT
AF287681.1  253 CCGATCG--TGCGTGTC-------------------ACCCCCACCGGCTTTGCGACTCT
GU572164.1  257 TCACGCG--CGACTCTGTCTGATTTGGACTACCGGACCCCTTCGACGTCTCC--------
MG236967.1  232 CCAAGCA--CGCGTGCTCCTGTTCG------GTGTGACCCTT---GGACCCC--------
AF202376.1  249 TCGTCCC--CGTCGTCTCGGGATGAGTCCTCAAGAGACCCTGTGCGAATGCG--------
MG234931.1  258 TCGTGTG--CATTTGTTGCTCG---------GGATGATCCTCAAACGAACCC--------
FJ428091.1  254 TCGTGCC--TTCGGGACCC------------GGGCGAGGCCTCGAGGACCCA--------
KX873060.1  249 CAGTGCA-------TCTGGCGCGTAGCCGACGTGATGGCCTTGAAGGACCCT--------
FN434493.1  246 CAGTGCA-------ACCGGCGCGCAGCCGGCGCAACGGCCCCAAAGGACCCA--------
JQ730037.1  285 TCGTG----GGCTCGGCGCCGGGC-------TGCGGGCTCCGCACAAGGACA--------
KF196362.1  275 GTGTGAA--TGTTTGTCACTT----------GGAAAGTTCTCTACCGACACC--------
AM492068.1  273 TCGTGCA--CGCTCGACCGCACACGGGGCCCCGGAGACCCCG--AAGAACCG--------
JN999661.1  252 CGGGACG--TGAACG----------------CAGAGGACTCG-TAGGACCCT--------
EU586943.1  239 TCGTGTG--CATGTATGCAGCCTTAGGATT-AGTAGACCCCTTTGCGTCCTT--------
KF677333.1  243 TCGTCCA--AATGTCGTGACGA---------GAAGGGCCCCT---GGATCCTTATGCGCA
KP098091.1  264 TTGAGGA--GCCGTGACGGCCCCCG------GGGCGCCCCCTCGAAAGACCC--------
KF498066.1  291 TCGTGTA--GGACTGCATTCGTT--------CCACGGGCACGACCTGATGGC--------
MF446079.1  271 TCGTGTG--GGAATGCG--------------TCGAGCGACGGGCACGACCCA--------
HE586032.1  240 TCACGCCTCTCCTTGCC--------------ACTGAGTTCTCGAAGGACCCT--------
KJ845982.1  274 CCACGTG--CGTCGGACACGCATCGGGACTCCGATAGACCCCAATGCGCCCG--------
KM064970.1  269 TCGCGCG--TGCCGCTCATCGCG--------CG------CTCCGCGAATCT---------
JN253228.1  253 TCGCGCG--CGTTCGAC--------------GGTCGAGACCC--TTGAACCC--------
KY858482.1  269 TTGTGAG--ATACAATTGTCGCAT-------TGTGGACATTATAATTGACCC--------
AY625633.1  264 CCGGGCT--CACTCGTTGCG-----------CGAGGGTACCT--CTGACCCT--------
KX646449.1  248 TCGGCCG--CTCTTGT---------------CTGGAAGCTCTTGATGACCCA--------
MG236571.1  232 CCGGTCG--CTCCTGT---------------CCATAAGCTCTCGTTGACCCA--------
AY936715.1  256 CCTCGTT--CATTTGTCCGAACCAA------CAAGGATCTCG--ACGACCCT--------
MG219588.1  241 TTGTGCG--TTACCGTCCTCTT---------CGTCGGCTCCA--CTGACCCT--------
AY613151.1  256 TCGTGT---CGTATTCCGTTGCATG------CTAGGGCATCTCTGCGACCCA--------
MG219618.1  237 TCGTGCG--TCCTAAGCTGTGAG--------GGATGTGCTCGATAAAGACCC--------
MG519650.1  260 CTGTGTG-TCGTGAGCTGCTAG---------GGAAGCCCTCAAAAAAGACCC--------
AB446176.1  234 TCGTGCC--GCCGAGCCGTCCGTGGTGATCCGAACGATGACCCAACGGAGCT--------
KU288713.1  252 TTGTGCG--CTTTCGTCGCGCTC--------CGAGCGGCCCGCGACGCTCGC--------
KM037612.1  251 TCGTGTG--CGCGTGTC--------------GAGCGAGGGCTCAACAAACCA--------
AY634813.1  250 TCATGCG--CGCGCGTCTGTC----------GGGGCCCCTCATCACGCACGT--------
EU528080.1  263 TCACGCG--ATCCTCCTTCGATTG-------AGGAGACTCCC--ACGACCCT--------
JQ255431.1  253 TCGCGGC--CACAGCCCGTCACGCT------CTCGGACTCCC---AGACCCT--------
AY233319.1  263 TCGTGCGGTGACCCGTCGCCAGCAAAAGATCTCATGACCCTGTTGCGCCGTC--------
```

```
AJ293780.1  303 ------ATTATGCGTGTGCAGGACGCCTTTTGGCTTT----------------CGACCTG
HQ246426.1  352 ---------------------TCTGCTTTCACCGCTGCCGCCGTCTAACCATTCGACCTG
KC411873.1  291 -------------------------TTT-----ACAC----------------TGACCTG
MF034641.1  267 ------------------CAGGAAATTCTCACCTTTT----------------CGACCTG
JN631369.1  360 -----------------------CTCTCTCAAAGCTT----------------GGACCTC
KP256679.1  368 TCGCGCATGGGAGGGGTGATGCTCCCTCTTC--GCAG----------------TGACCCC
KX229790.1  339 ---CGCCTTGGCAATGCCACGCGCGTCGCCCGGACAC----------------CGGCCCC
GQ866430.1  339 -CCGATCAATGCGTCGTATGTGACACCTTGG--AATG----------------CGACCCC
AJ489792.1  297 -------TGGATTGAGCATCGATTGCTTGC---ACCG----------------CGACCCC
KT825439.1  300 ----------------------ATTTTCCTTTGAAGG----------------TGATCCC
KP057199.1  300 ----------------------ATTTTCCTTCGAAGG----------------TGATCC-
FJ213861.1  305 --AGAATAGCACATGCCGAATGGCATTGGCGAAACCA----------------AGACCCC
KX087731.1  249 ATTGTGTTGTTTGTCTGTAATAATGATGTACTTATCG----------------AGACCTC
AF287681.1  291 ACGGCCCATGAGCGTCTTTTGGTCGCCCAA---GACG----------------GGACCTC
GU572164.1  307 --------------------GGACGCTCTTT--AGCG----------------AGACCTC
MG236967.1  273 ---------TGCGGCGTCTTCGGATGCTCTTG----CG----------------AGACCTC
AF202376.1  299 -----GCGTCGGCGAAAGTGCCGCGCCCATCAAATTG----------------TGGCCCC
MG234931.1  299 -----CTACACGTCTTAAATCGACGCTATT---GTCG----------------CGACCCC
FJ428091.1  292 -----AGTCGTGGTGCGAGTCGATGCCACGG--ACCG----------------CGACCCC
KX873060.1  294 ------ATGAACGGTGCGCACGACGCTCCG---ACCG----------------CGACCCC
FN434493.1  291 ------ACGAACGGAGCGCACGTCGCTCCG---ACCG----------------CGACCCC
JQ730037.1  326 ------------------------ATCCCCA--AGTG----------------CGATCCC
KF196362.1  315 -----TTGAACGTCTTTTAATGACCGTTCTACTGTCG----------------CGACCCC
AM492068.1  321 ----------TCGTCAAGGACGACGCTCTC---ACTG----------------CGACCCC
JN999661.1  285 -------TAGCTGTCTCCTTTGGCGACAGAACCGTTG----------------CGACCCC
EU586943.1  288 ----------------TGTGGCACGCTCAC---ATTG----------------CGACCCC
KF677333.1  289 TTAGTCACTCACTGAGGGATGATTGCTCACG--ACCG----------------CGACCCC
KP098091.1  308 ----------CATTGAGAGCAGTGGCCTCCG--ACCG----------------CGACCCC
KF498066.1  333 ---------------AGAAGGCTTGCCCTCG--ATTG----------------CGACCCC
MF446079.1  307 ----------ACGGCAGCAAGATTGCTCTCG--TTCG----------------CGACCCC
HE586032.1  278 ---TGTGTGTGTTGCCTCGTTGCAACCCTCACCGTTG----------------CGACCCC
KJ845982.1  324 ----------------TTGCGGGTGCCTCC---AACG----------------CGACCCC
KM064970.1  304 ---------------------GCTCCTTACC--AACG----------------CGACCCC
JN253228.1  287 ---TTTCGGCATCGCAAGGACGGTGCTCGC---ATCG----------------CGACCCC
KY858482.1  312 -----------AACAATGCCAAGTGCTTTCG--ACCG----------------CGACCCC
AY625633.1  301 ---ATGTGCGTCCCTTGGGGAATTGCTCGCA----GG----------------CGACCCC
KX646449.1  283 ----------------------AAGTCCTC---AACG----------------CGACCCC
MG236571.1  267 ----------------------ATGTCATC---AAAG----------------CGACCCC
AY936715.1  298 --------------------CTATGTATCC---GACG----------------CGACCCC
MG219588.1  280 -----GTTGCATTGCGAAAGTGATGCCT-CG--ACCG----------------CGACCCC
AY613151.1  299 ---------ACGGCACACACGTGTGCTTCCG--ACCG----------------AGACCCC
MG219618.1  279 --CAATGTGTCGTCCTGCGACGATGCTTCG---ACCG----------------CGACCCC
MG519650.1  302 ---CATTGTATCGTCTTAGATGATGCTTCG---ACCG----------------CGACCCC
AB446176.1  284 -----------------------TGCTCCTTCGACCG----------------CGACCCC
KU288713.1  294 ------TGCTCTCGCTTCTGCGGAGCTTTC---AACG----------------CGACCCC
KM037612.1  287 ----TGTTGCATCGATTCGTCGATGCTTTC---AACG----------------CGACCCC
AY634813.1  290 ------------CTACCCGTCGACGCTCTC---AACG----------------CGACCCC
EU528080.1  304 ---GATGCATGCGTTGTCATGACGCCTGCCACGACCG----------------CGACCCC
JQ255431.1  294 ---------GTTCGCGATTATGGTGCTCCG---ACCA----------------CGACCCC
AY233319.1  315 --------------CTCGACGCGCGCTCCG---ACCG----------------CGACCCC
```

```
AJ293780.1  341 AGGTCAGGCAAGAACACCC-------------------------------
HQ246426.1  391 AGCTCAGGCAAGATTACCCGCTGA--------------------------
KC411873.1  305 AGATCAGACAAGGCTACCCGCTGAACTTAAGCAT----------------
MF034641.1  293 AGATCAGGCAAG--------------------------------------
JN631369.1  381 AGATCAGGCAAGATTACCCGCTGAG-TTAAGCATATCACTAAGC------
KP256679.1  410 AGTTCAGGCGAGGGTACCCGCCGAGTTTAAGCATATAACTAAGCGGAGGA
KX229790.1  380 AGGTCAGACGGGAGCACCCGCT----------------------------
GQ866430.1  380 AGGATGGGCGGGA-------------------------------------
AJ489792.1  331 AGGT----------------------------------------------
KT825439.1  322 A-------------------------------------------------
KP057199.1      --------------------------------------------------
FJ213861.1  347 AGGTCAGG------------------------------------------
KX087731.1  293 AGGTCAGGCGGGGCTACCCGCTGAGTTTAAGCATATCAA-----------
AF287681.1  332 AGGTCAGGCGGGGCTACCCGCTGAGTTTAAGCA-----------------
GU572164.1  329 AGGTCA--------------------------------------------
MG236967.1  305 AGGTCAGGCGGGGCTACCCGCTGAGTTTAA--------------------
AF202376.1  338 AAGTCAGGCGAG-CCACCCGCCGA--------------------------
MG234931.1  335 AGGTCAGGCGGGATTACCCGCTGAGTTTAA--------------------
FJ428091.1  329 GGGTCAGGTGGGGCTACCCGCTGAG-------------------------
KX873060.1  329 AGGTCAGGCGGGACTACCCGCTGAATTTAAGCATATAAATAAGCGG----
FN434493.1  326 AGGTCA--------------------------------------------
JQ730037.1  344 AGGTCAGGCGGGGCTACCCGCTGAGTTTAAGCATATCAATAAGCGGAGGA
KF196362.1  354 AGGTCAGGCGGGATTACCCGCTGAATTTAAGCATATCAATAAGCGGAGGA
AM492068.1  352 AGGTCAGGCGGGGCCACCCGCTGAGTT-----------------------
JN999661.1  322 AGGTCAGGCGGGGCTACCCGCTGAGTTTAA--------------------
EU586943.1  313 AGGTCAGGTGGGATTACCCGCTGAGTTTAAGC------------------
KF677333.1  331 AGGTCAGGC-----------------------------------------
KP098091.1  340 AGGTCAGGC-----------------------------------------
KF498066.1  360 AGGTCAGGCGGGATCACCCGCTGAGTTTAAGCATATCAATAAGCGGAGGA
MF446079.1  339 AG------------------------------------------------
HE586032.1  319 AGGTCAGGCGGGGCTACCCGCTGAGTTTAAGCATATC-------------
KJ845982.1  349 AGGTCAGGCGGGGCTACCCGCTGAGTTTAAGCATATCAA-----------
KM064970.1  325 AGGTCAAGCGGGGCTACCCGCTGAGTTTAAGCATATCA------------
JN253228.1  325 AGGTCAGGCGGGATTACCCGCTGAGTTTAA--------------------
KY858482.1  343 AGGTCAGGCGGGATTACCCGCTGAGTTTAAGCATATCAATAAGCGGAGGA
AY625633.1  338 AGGTCAGGCG----------------------------------------
KX646449.1  302 AGGTCAGGCGAGATCACCCGCTGAGTTTAAGCATATCAAT----------
MG236571.1  286 AGGTCAGGCGGGATCACCCGCTGAGTTTAAGCATATCAATAAGCGGAGG-
AY936715.1  319 AGGTCAGGCGGGATTACCCGCTGAGTTTAAGCATATCAATAAGCGGAGGA
MG219588.1  316 AGGTCAGGCGGGACTACCCGCTGAGTTTAA--------------------
AY613151.1  332 ATGTCAGGCG-GATTACCCG------------------------------
MG219618.1  318 AGGTCAGGCGGGACTACCCGCTGAGTTTAA--------------------
MG519650.1  340 AGGTCAGGCGGGACTACCCGCTGAGTTTAAGCATATCATAAAGC------
AB446176.1  305 AGTGCAGG------------------------------------------
KU288713.1  329 AGGTCAGGCGGG--------------------------------------
KM037612.1  324 AGGTCAGGCGGGGTTACCCGCTGAATTTAAGCATATCAATAAGCGGAGGA
AY634813.1  319 AGGTCAGGCGGGGTTACCCGCTGAATTTAA--------------------
EU528080.1  345 AGGTCAGGCGGGATTACCCGCT----------------------------
JQ255431.1  326 AGGTCAGGCGGGACTACC--------------------------------
AY233319.1  342 AGGTCAGGCGGGACTACCCGCTGAGTTTAAGCATATCAATAAGCGGAGGA
```

**References**

Taberlet P., Gielly L., Pautou G. and Bouvet J. (1991) Universal primers for amplification of three non-coding regions of chloroplast DNA. *Plant Molecular Biology*, **17**, 1105-1109.

Chen S., Yao H., Han J., et al. (2010). Validation of the ITS2 region as a novel DNA barcode for identifying medicinal plant species. *PLoS ONE*, **5**, e8613.

White T. J., Bruns T., Lee S. and Taylor J. W. (1990). Amplification and direct sequencing of fungal ribosomal RNA genes for phylogenetics. *In* M. A. Innis, D. H. Gelfand, J. J. Sninsky, & T. J. White (Eds.), PCR pro- tocols: A guide to methods and applications (pp. 315–322). New York, NY: Academic Press.

Kress W.J. and Erickson D.L. (2007) A two-locus global DNA barcode for land plants: The coding rbcL gene complements the non-coding *trnH-psbA* spacer region. *PLoS ONE*, **2**, e508.

Palmieri L., Bozza E. and Giongo L. (2009) Soft fruit traceability in food matrices using real-time PCR. *Nutrients*, **1**, 316-328.

Elbrecht V. and Leese F. (2017) Validation and development of COI metabarcoding primers for freshwater macroinvertebrate bioassessment. *Frontiers in Environmental Science*, **5**, 11.
